# Celsr1 and CAMSAP3 differently regulate intercellular and intracellular cilia orientation in oviduct multiciliated cells

**DOI:** 10.1101/2020.08.28.273169

**Authors:** Fumiko Matsukawa Usami, Masaki Arata, Dongbo Shi, Sanae Oka, Yoko Higuchi, Fadel Tissir, Masatoshi Takeichi, Toshihiko Fujimori

## Abstract

The molecular mechanisms by which cilia orientation is coordinated within and between multiciliated cells (MCCs) is not fully understood. By observing the orientation of basal bodies (BB) in MCCs of mouse oviducts, here, we show that Celsr1, a planar cell polarity (PCP) factor involved in tissue polarity regulation, is dispensable for determining BB orientation in individual cells, whereas CAMSAP3, a microtubule minus-end regulator, is critical for this process but not for PCP. MCCs exhibit a characteristic BB orientation and microtubule gradient along the tissue axis, and these intracellular polarities were maintained in the cells lacking Celsr1, although the intercellular coordination of the polarities was partly disrupted. On the other hand, CAMSAP3 regulated the assembly of microtubules interconnecting BBs by localizing at the BBs, and its mutation led to disruption of intracellular coordination of BB orientation, but not affecting PCP factor localization. Thus, both Celsr1 and CAMSAP3 are responsible for BB orientation but in distinct ways; and therefore, their cooperation should be critical for generating functional multiciliated tissues.

## Introduction

Multiciliated cells (MCCs), which have a few hundred cilia on their apical surface (Brooks and Wallingford, 2014; Meunier and Azimzadeh, 2016; Reiter and Leroux, 2017; Spassky and Meunier, 2017; Boutin and Kodjabachian, 2019), are specialized cells found in the epithelium of the trachea and the ependyma of the brain. In the infundibulum of the mouse oviduct (Fallopian tube), eggs released from the ovaries are transported to the uterine side by the movement of multicilia (Li and Winuthayanon, 2016; Shi et al., 2011). In MCCs, the direction of cilia movement is coordinated both intracellulary (within each cell : rotational polarity) and intercelluary (between cells tissue-level polarity) along the axis of the organ to generate directed flow(Mirzadeh et al., 2010; Meunier and Azimzadeh, 2016; Boutin et al., 2014; Tissir et al., 2010b; Hirota et al., 2010; Guirao et al., 2010). Ciliary abnormalities, including motility and polarity as well as ciliogenesis in various organs, can lead to various disorders (Choksi et al., 2014; Reiter and Leroux, 2017).

This unidirectional ciliary movement is reinforced by the structural polarity of each cilium. Motile cilia exhibit a 9+2 axonemal microtubule organization, and the alignment of the central pair of microtubules is perpendicular to the direction of the effective strokes (Sandoz et al., 1988). The basal body (BB) found at the base of the cilium has similar components to the centriole and supports the axoneme in the apical cytoplasm. The basal foot (BF) (Reiter et al., 2012; Clare et al., 2014; Garcia and Reiter, 2016), an electron-dense structure that projects unidirectionally from the BB in the same orientation as the effective stroke.

The orientation of multiple cilia is reflecting the polarity of each cell. In epithelial cells, in addition to apicobasal polarity, planar cell polarity (PCP) runs perpendicular to the apicobasal plane within the plane of the epithelium (Butler and Wallingford, 2017; Henderson et al., 2018; Aw and Devenport, 2017). For example, in the oviduct, the axis of the epithelial PCP lies along the longitudinal axis of the oviduct, and the morphological polarities extend over multiple cells, such as the folded structure formed by the epithelial sheet, the elongation of the apical surface of MCCs, and the orientation of multicilia (Shi et al., 2014; Koyama et al., 2016; Koyama et al., 2019). PCP factors are proteins involved in regulating PCP that were initially identified by genetic screens performed in *Drosophila melanogaster*; many are conserved in vertebrates (Butler and Wallingford, 2017; Devenport 2014; Yang and Mlodzik, 2015; Hale and Strutt, 2015; Boutin et al., 2012; Shi et al., 2013; Shi et al., 2016a). These factors localize asymmetrically in each cell according to the tissue’s polarity. For example, in the mouse oviduct, Vangl1 (Van Gogh-like protein 1) and Vangl2 are localized to the cell boundary on the ovarian side, while Fzd (Frizzled) 6 is localized to the uterine side (Shi et al., 2014; Shi et al., 2016b). Furthermore, Celsr1 (Cadherin EGF LAG seven-pass G-type receptor 1), a seven-transmembrane protein with extracellular cadherin repeats, localizes to the cell boundary of both the ovarian and uterine sides. Dvl (Dishevelled), a cytoplasmic PCP factor, also localizes to the base of cilia, in addition to cell boundaries in *Xenopus laevis* epidermis and mouse ependyma (Hirota et al., 2010; Ohata et al., 2014; Park et al., 2008; Mitchell et al., 2009). PCP factors are essential for regulating cilia polarity in *Xenopus* epidermis (Mitchell et al., 2009; Park et al., 2008), mouse ventricular ependymal cells (Tissir et al., 2010a; Hirota et al., 2010; Guirao et al., 2010; Mirzadeh et al., 2010; Boutin et al., 2014; Ohata et al., 2014; Takagishi et al., 2017; Takagishi et al., 2020), mouse trachea (Vladar et al., 2012), and mouse oviduct (Shi et al., 2014). We previously reported that Celsr1 deficient females are sterile because of multiple PCP defects in the mutant oviducts (Shi et al., 2014). By using fluid flow analysis and direct observation of ciliary movement, we found that the directions in which the cilia of MCCs move were not aligned along the longitudinal axis of the oviduct. Our analysis following electron microscopy observation has revealed that cilia orientation is not coordinated in *Celsr1-/-* mutant oviducts. But we could only observe details in a very limited area, and could not simultaneously determine BB orientation across multiple cells. Although the contribution of PCP factors to the coordination of cilia orientation is evident, the mechanisms by which PCP factors communicate with cilia remain unclear.

Cytoskeletal elements, namely microtubules, intermediate filaments, and actin filaments are connected to the BB (Sandoz et al., 1988; Werner et al., 2011; Clare et al., 2014; Antoniades et al., 2014; Chevalier et al., 2015; Herawati et al., 2016; Tateishi et al., 2017), and supposedly provide mechanical support for cilia. The mechanism by which these cytoskeletal elements are involved in cilia polarity regulation differs between animal species and between organs. In *Xenopus* epidermal cells, actin filaments play a major role in regulating cilia polarity, positioning, and motility, and microtubules contribute to regulating local polarity between adjacent cilia (Werner et al., 2011; Antoniades et al., 2014). Actin filaments extending from neighboring BBs are bound to the rootlets. Apart from these structures, microtubules form a network connecting the BBs. Cilia are connected via microtubules (Iftode and Fleury-Aubusson, 2003; Bayless et al., 2019; Soh et al., 2019) and aligned to coordinate cilia orientation in ciliophora (ciliates). In mouse trachea, actin filaments function in BB docking to the apical surface of cells, and microtubules regulate cilia orientation (Herawati et al., 2016). Electron microscopy has revealed that microtubules accumulate near the tip of the BF in various animal species (Reed et al., 1984; Lemullois and Marty, 1990; Sandoz et al., 1988). In previous reports, MCCs were treated with drugs that affect either microtubule polymerization or depolymerization in order to determine whether microtubules regulate cilia orientation. However, microtubules were organized and distributed within MCCs in a variety of ways all of which could be disrupted by drugs. Therefore, more careful studies are required to determine which microtubules specifically coordinate cilia orientation. Moreover, the molecular mechanisms underlying microtubule control of cilia orientation remain unclear.

In this study, we explored how the orientation of cilia is coordinated within each cell and between cells, focusing on MCCs in the infundibulum of the mouse oviduct. To this end, we made use of super-resolution microscopy to dissect the roles of PCP factors and microtubules in establishing tissue-wide coordination of cilia orientation. In addition to the role of Celsr1, a PCP factor that organizes the orientation of cilia and polarity of microtubule enrichment at the tissue level, we found that CAMSAP3, a microtubule minus end-binding protein, localizes to BBs, controls rotational polarity of cilia, and organizes the assembly of microtubules interconnecting BBs.

## Results

### Intracellular basal body orientation is moderately coordinated in *Celsr1*-deficient oviduct MCCs

To investigate the orientation of individual cilia across an epithelial sheet, we used super-resolution microscopy. To determine cilia orientation, oviducts were stained with anti-Odf2 antibody (Tateishi et al., 2013), a BB (distal and subdistal appendage) marker, and anti-Centriolin antibody (Mazo et al., 2016), a BF marker. Odf2 formed a ring with an approximate diameter of 280 nm, and Centriolin was detected as an ellipse with a major axis of about 160 nm, or as two adjacent puncta (Fig. 1A). The orientation of each BB was determined by the relative positions of Odf2 and Centriolin (Fig. 1A’ and A’’), which corresponds to the direction of the ciliary effective stroke. The average number of BBs in adult wild-type (WT) MCC was 201 per cell (n = 20 cells, SD = 25.7). We determined the orientation of 183 BBs in each cell on average (n = 20 cells; Fig. 1B and C), choosing those with clear Odf2 and Centriolin signals. The variation in BB orientation within each cell is indicated as cellular circular variance (CV) (Fig. 1D) (Shi et al., 2014), where cells with lower CV represent uniform BB orientation. Centriolin signals were localized to the uterine side of the Odf2 ring (Fig. 1B), indicating that BBs were regularly oriented in the same direction in each cell. The CVs were less than 0.2 (median = 0.098) in all the cells, suggesting uniform cilia orientation in WT cells. We also calculated the cellular mean BB orientation in each cell (Fig. 1C) and compared this value between adjacent cells (Fig. 1E). The mean BB orientation was also uniform between cells heading to the uterine side in the WT.

**Figure 1.**
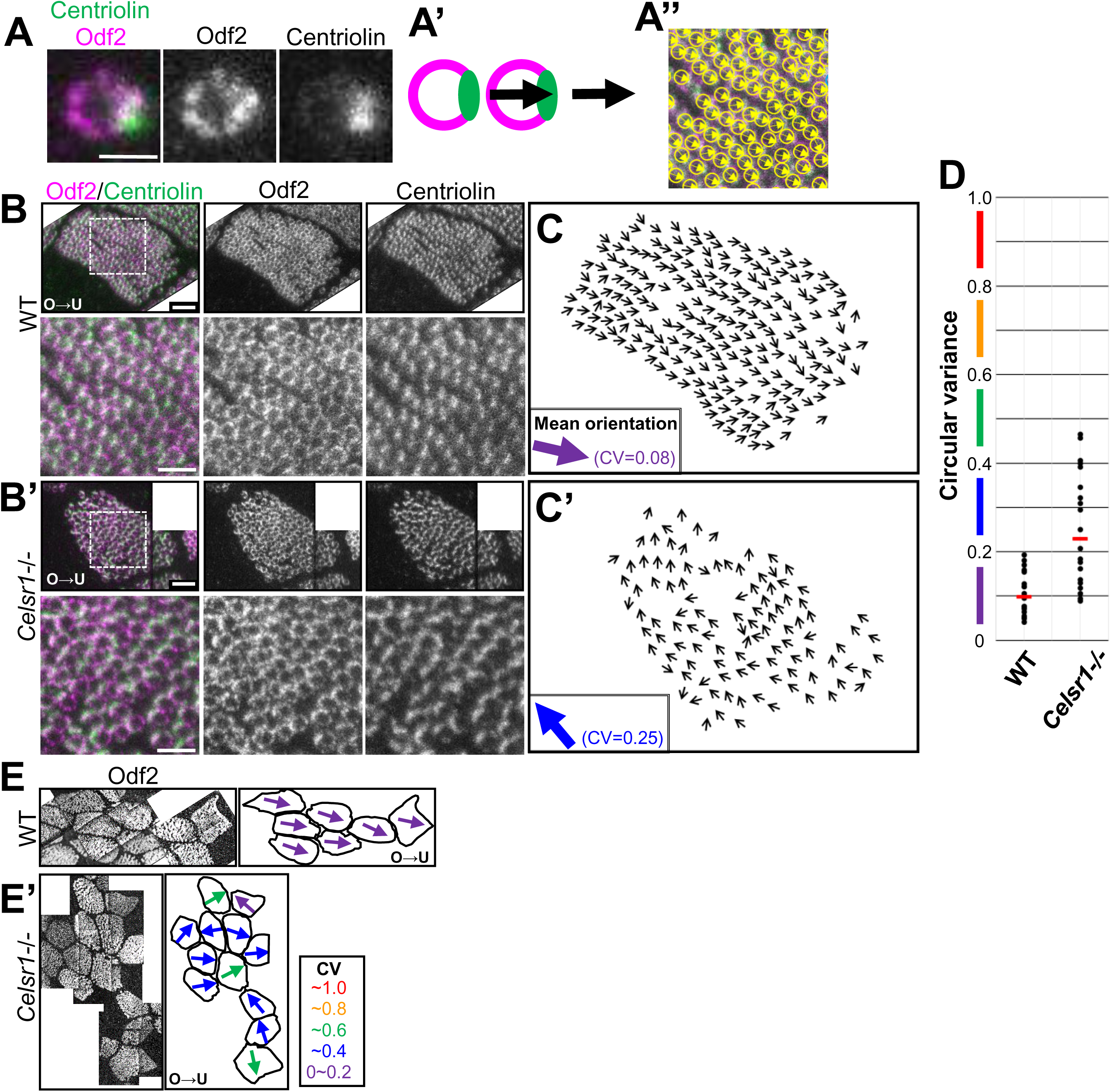
Basal body orientation in *Celsr1* deficient oviduct multi-ciliated cells (MCCs) A, Super-resolution microscope (STED) image of a basal body (BB) of WT oviduct MCCs after staining with anti-Odf2 (magenta) and anti-Centriolin (green) antibodies. (Bar = 0.3 μm) A’ and A’’, On the Odf2 ring, the arrow points toward the center of the Centriolin signals and corresponds with BB orientation. A’’, An example showing the determination of BB orientation in an actual image (same image as B, bottom). B, 12-week-old (12 w old) WT MCC stained with anti-Odf2 (magenta) and anti-Centriolin (green) antibodies. Images were manually tiled because the cell could not be captured in one field, and maximum intensity projection (MIP) images (rendered using up to 15 slices acquired at a step size of 0.1 μm to cover all Centriolin and Odf2 signals) are shown. The top row shows the original image (Bar = 1 μm) while the bottom row shows the high magnification image (Bar = 0.5 μm). In this cell, 210 BBs were analyzed and the CV was 0.08. The ovarian and uterine sides are on the left and the right respectively in all figures (shown as O > U). B’, *Celsr1-/-* mutant, 121 BBs were analyzed (CV = 0.25). C and C’, Small arrows indicate BB orientation in cells shown in B and B’. The mean BB orientation is indicated by the large arrow, and the arrow color represents the CV value of the cell. D, The variation in the orientation of BBs in each cell was evaluated as CV. CV values range from 0 (totally unidirectional) to 1 (no bias). Each point on the graph corresponds to one cell and the red bar represents the median. The colors are used to indicate the CV value of each cell when showing the mean BB orientation of the cell with the arrow, as shown in Fig. 1C, C’ and E, E’. Twenty cells from two animals were analyzed for both genotypes. The datasets are identical to those shown in Figure S5. E and E’, Representatives of mean BB orientation in neighboring WT cells and *Celsr1-/-* cells, respectively. Tiled images were used to compare angles between cells. The mean BB orientation of each cell is shown with an arrow, and the arrow colors represent the CV values shown in the bottom right inset.

In *Celsr1-/-* mutant, both Odf2 and Centriolin signals were detected in each BB with the same shape and in the same relative positions as in WT. The average number of BBs in adult *Celsr1-/-* mutant cells was 199 per cell (n = 20 cells, SD = 27.7), which was comparable with WT. We then analyzed the orientation of 127 BBs per cell on average (n = 20 cells; Fig. 1B’ and C’). The variation in BB orientation within each *Celsr1-/-* mutant cell was compared with WT. The CV values for the *Celsr1-/-* mutants were between 0.09 and 0.5 (median = 0.23), which was significantly higher than WT, but still lower than completely random (Fig. 1D and Fig. S5A, A’). These results are consistent with our previous study using electron microscopy (Shi et al., 2014). In the *Celsr1-/-* mutant, the mean BB orientation varied between cells. This was the case even when BB orientation is approximately aligned with lower CV values in neighboring cells (Fig. 1E’, purple and blue arrows).

These analyses confirmed that BBs in each cell are not randomly oriented in WT, and *Celsr1-/-* mutants show reduced coordinated orientation of BBs, and the mean BB orientation differed between cells. These results are consistent with our previous finding that fluorescent beads were not transported in a single direction in *Celsr1-/-* oviducts (Shi et al. 2014). This consistency suggests that Celsr1 is involved in coordinating cilia orientation between cells, although other unidentified mechanisms appear to exist for aligning cilia orientation in each cell even when Celsr1 is absent.

### Polarized microtubule enrichment at the cell-cell boundary correlates with BB orientation in each cell even in the absence of Celsr1

The microtubule network has been proposed to support cilia orientation in mammalian cells (Herawati et al., 2016; Vladar et al., 2012). In addition, microtubules are concentrated at the apical cell-cell boundary in a polarized manner in MCCs of mouse ependyma and trachea; this polarized enrichment correlates with the orientation of BBs (Boutin et al., 2014; Herawati et al., 2016; Takagishi et al., 2017; Vladar et al., 2012). We have also previously shown that EB1, which localizes to the plus-end of microtubules, is enriched in a polarized manner in mouse oviduct MCCs (Shi et al., 2016b).

We then examined the relationship between the organization of microtubules and BB orientation in oviduct MCCs of WT and *Celsr1-/-.* In the WT, microtubules visualized by anti–β-tubulin antibody staining were asymmetrically enriched on the uterine side of the cell-cell boundary in each MCC (Fig. 2A). Moreover, a gradient of signals to this enrichment was observed near the apical surface (Fig. 2A and Fig. S1A). Quantitative analyses clearly indicated microtubule enrichment on the uterine side of MCCs in WT (Fig. 2B). In contrast, in the *Celsr1-/-* mutant, asymmetric enrichment of microtubules was evident in >75% of MCCs, but the direction of microtubule enrichment did not match between cells, nor did it match either the ovarian-uterine axis or fold morphology (Fig. 2A’ arrows and Fig. 2B’).

**Figure 2.**
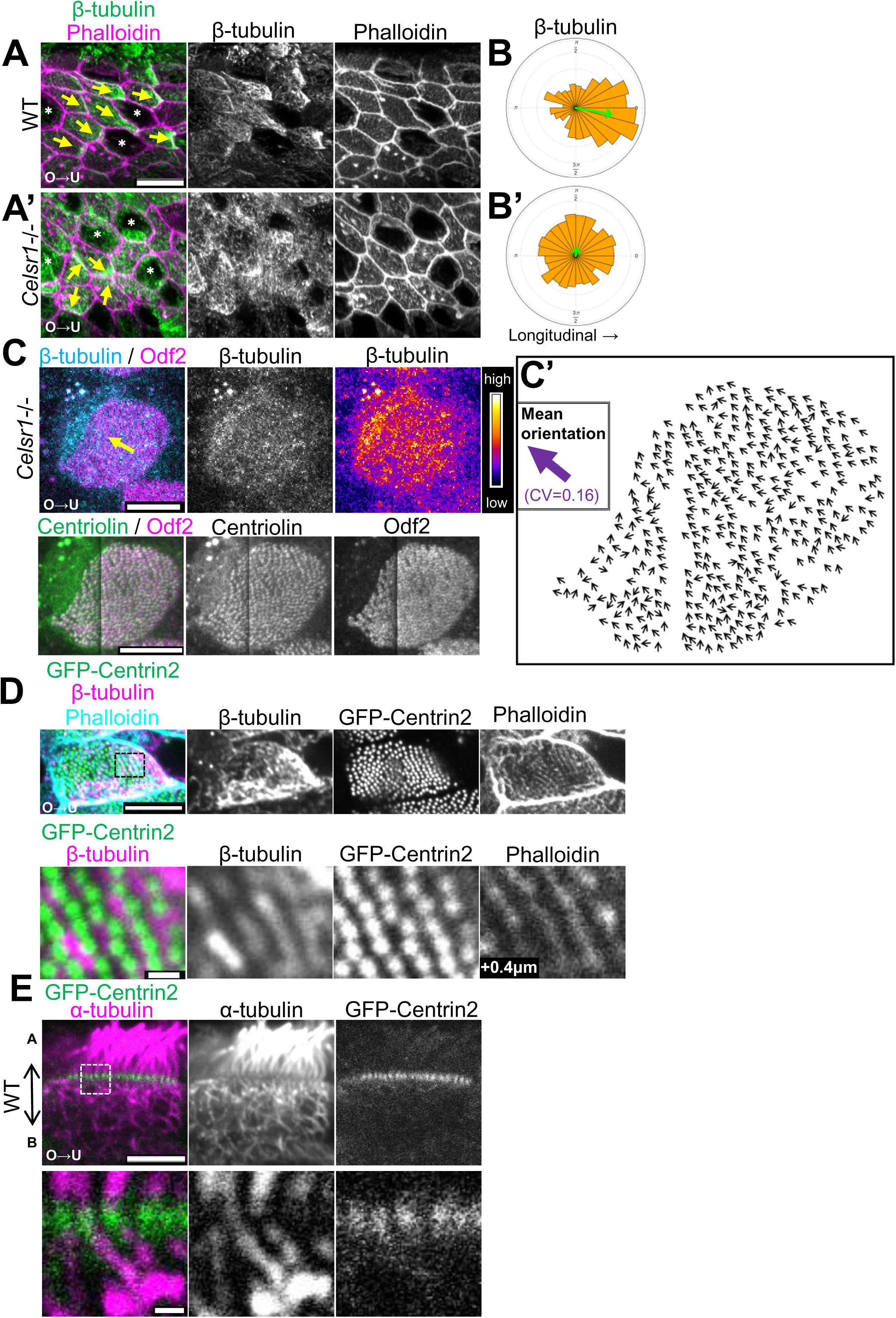
Direction of polarized microtubule enrichment correlated with BB orientation in WT and *Celsr1-/-*. A and A’, Oviduct epithelium of adult (11 w old) WT (A) and *Celsr1-/-* mutant (A’) stained with anti–β-tubulin (green) and phalloidin (magenta). MIP images created from 20 images acquired at a step size of 0.2 μm (Bar = 10 μm). The direction of β-tubulin enrichment is shown with yellow arrows. Secretory cells are indicated with white asterisks. B and B’, Rose diagrams show the results of quantifying the orientation of microtubule enrichment near the cell-cell boundary. The angles between the elongated folds (longitudinal axis of the folds) were measured. The angle was classified into 24 bins. The area of each bin is proportional to the number of cells within each bin. A total of 370 cells from three WT animals and 348 cells from three *Celsr1-/-* animals were analyzed. The angle and the length of each green arrow indicates the mean angle and 1 minus CV, respectively. C, Top row, confocal images of the *Celsr1-/-* mutant oviduct stained with anti–β-tubulin (blue) and anti-Odf2 (magenta) antibodies. MIP images were rendered from 10 images acquired at a step size of 0.3 μm. A yellow arrow indicates the orientation of microtubule enrichment. Bottom row, super-resolution microscopy images stained with anti-Odf2 (magenta) and anti-Centriolin (green) antibodies. MIP images were rendered from 20 images acquired at a step size of 0.1 μm. To obtain Centriolin signals, the fixation condition was different from that used for Fig. 2A and A’ (Bar = 5 μm). C’, Small arrows indicate BB orientation, and color represents angles from mean BB orientation as shown in the inset. D and E, Apical view (D) and lateral view (E) of MCCs of *GFP-CETN2* (green) transgenic mouse stained with anti–β-tubulin (magenta in D) or anti–α-tubulin (magenta in E) antibodies and phalloidin (cyan). D, Top row, MIP image was rendered from three images acquired at a step size of 0.2 μm. Bottom row shows a single plane at high magnification. Focal plane of phalloidin was 0.4 μm basal to GFP-Centrin2 and β-tubulin, and indicated as “+0.4 μm” in D. E, single plane images. Bottom row, high magnification after background subtraction. The apical side is located towards the top of each image. (Bar = 5 μm and 0.5 μm in the top and bottom rows respectively in D and E.)

We next analyzed whether there is any correlation between the orientation of microtubule enrichment and BB orientation in *Celsr1-/-*. BB orientation was determined by Odf2 and Centriolin signals as shown in Fig. 1. Since the optimal fixation conditions for the anti–β-tubulin antibody and for the anti-Odf2 and anti-Centriolin antibodies are different, the respective staining signals were slightly different from that which is described above (Fig. 1A and Fig. 2A’). The β-tubulin signal was observed in the form of fine dots, nevertheless allowing us to observe its density gradient (Fig. 2C). The BB orientation was determined and quantified in cells where microtubule enrichment was observed (Fig. 2C’). These cells showed CV values lower than 0.25 (median = 0.14), and many BBs were oriented in similar directions in each cell. The mean BB orientation (the big purple arrow in Fig. 2C’) in each cell coincided with the microtubule enrichment direction (the yellow arrow in Fig. 2C; n = 5 cells, average = 210 BBs/cell).

Thus, microtubules were enriched near the apical surface of MCCs even in the absence of Celsr1, and the orientation of microtubule enrichment correlated with the mean BB orientation in each cell; however, microtubule enrichment orientation between cells was not aligned. These results indicate that BBs are oriented towards the region of microtubule enrichment near the apical surface, and Celsr1 plays a role in aligning the orientation of microtubule enrichment between cells.

### Connection between microtubules and BBs in oviduct MCCs

Electron microscopic observation of MCCs has previously shown that microtubules are attached to the tip of the BF and run along the apical surface interconnecting adjacent BFs (Herawati et al., 2016; Vladar et al., 2012; Tateishi et al., 2017; Sandoz et al., 1988; Reed et al., 1984). It remains unclear, however, how the microtubules are connected to the BBs and how they are involved in organizing BB orientation. In order to determine microtubule localization and elucidate the relationship between microtubules and BB/BF in detail, microtubules and BBs were visualized by using an anti–β-tubulin antibody and GFP-Centrin2, respectively. Microtubules formed a striped pattern on the plane parallel to the apical surface in the range of 0.6 μm, which was identical to GFP-Centrin2 (Fig. 2D and Sup. Mov.1). In cells with aligned BBs, microtubules were also linearly aligned to form a striped pattern along the BBs, partly overlapping with the latter (Fig. 2D bottom). The signal intensities of the stripes were also stronger on the uterine side of each cell (Fig. 2D) and weaker on the ovarian side. When examined from the lateral side of the cells, a thick microtubule bundle (visualized with anti–α-tubulin antibody) connected to each BB (visualized with GFP-Centrin2) extended to the basal side of the cell in the cytoplasm (Fig. 2E), although which components of microtubules visualized by lateral sections correspond to the apical microtubule stripes, shown in Fig. 2D, remains unclear. We also examined *Celsr1-/-* oviducts and found that the apical microtubule stripes were maintained while apico-basal bundles were clearly connected to BBs (Fig. S1A and B). To sum, oviduct MCCs have microtubules running along BBs as well as those extending basally from BBs, and either one of them or both are enriched at a cell boundary; and Celsr1 is not involved in the assembly of these microtubules, though it is important for their PCP orientation.

### CAMSAP3, a microtubule minus-end binding protein, localizes asymmetrically to the base of cilia

Both apical microtubules forming stripes and microtubule bundles running apico-basally appear to connect to BBs, and these might be differentially involved in the regulation of cilia orientation. Although γ-tubulin localizes in BBs and BFs (Muresan et al., 1993; Hagiwara et al., 2000), it is unknown if γ-tubulin regulates ciliary orientation (Yuba-Kubo et al., 2005). The minus-ends of axonemal microtubules of the cilium are oriented towards the BBs in MCCs. In intestinal non-ciliated epithelial cells, the minus-ends of non-centrosomal microtubules that run apico-basally are anchored to the apical surface by CAMSAP3 (calmodulin-regulated spectrin-associated protein3), or Nezha (Meng et al., 2008), a non-centrosomal microtubule minus-end binding protein (Toya et al., 2016). Based on the above, we sought to determine if CAMSAP3 plays a role in MCCs and is involved in organizing microtubules to regulate the orientation of BBs.

We compared CAMSAP3 distribution in MCCs with that of other BB/BF components. CAMSAP3 was detected on the uterine side of BBs labeled with GFP-Centrin2, and these two signals were observed as pairs (Fig. 3A). When observed from the lateral side of the cells, CAMSAP3 partially overlapped GFP-Centrin2 on the apical side (Fig. 3A’). γ-tubulin localized to the uterine and basal sides of the GFP-Centrin2 signal, and these two signals were also paired (Fig. 3B and B’). When CAMSAP3 and γ-tubulin were compared directly, both signals were detected as small dots in the apical view (Fig. 3C). In the lateral view, these two signals were separated, and the CAMSAP3 signal showed apical localization relative to the γ-tubulin signal (Fig. 3C’). CAMSAP3 and γ-tubulin seemed to be paired, but the positional relationship between them was not constant in the *xy*-plane. This may be due to the longer distances between the two signals along the apico-basal axis, and the BB and axoneme were not always vertical relative to the apical surface. Hence, the angles between these were variable (Fig. S2B, bottom panels, dotted lines) when MCCs were observed from the lateral side. CAMSAP3 was detected as a small dot on the same side as Centriolin, which was found to be located away from the center of BBs using super resolution microscopy (Fig. 3D). CAMSAP3 also showed apical localization relative to Centriolin. We also observed CAMSAP3 signals on the uterine side of Odf2 (Fig. S2C).

**Figure 3.**
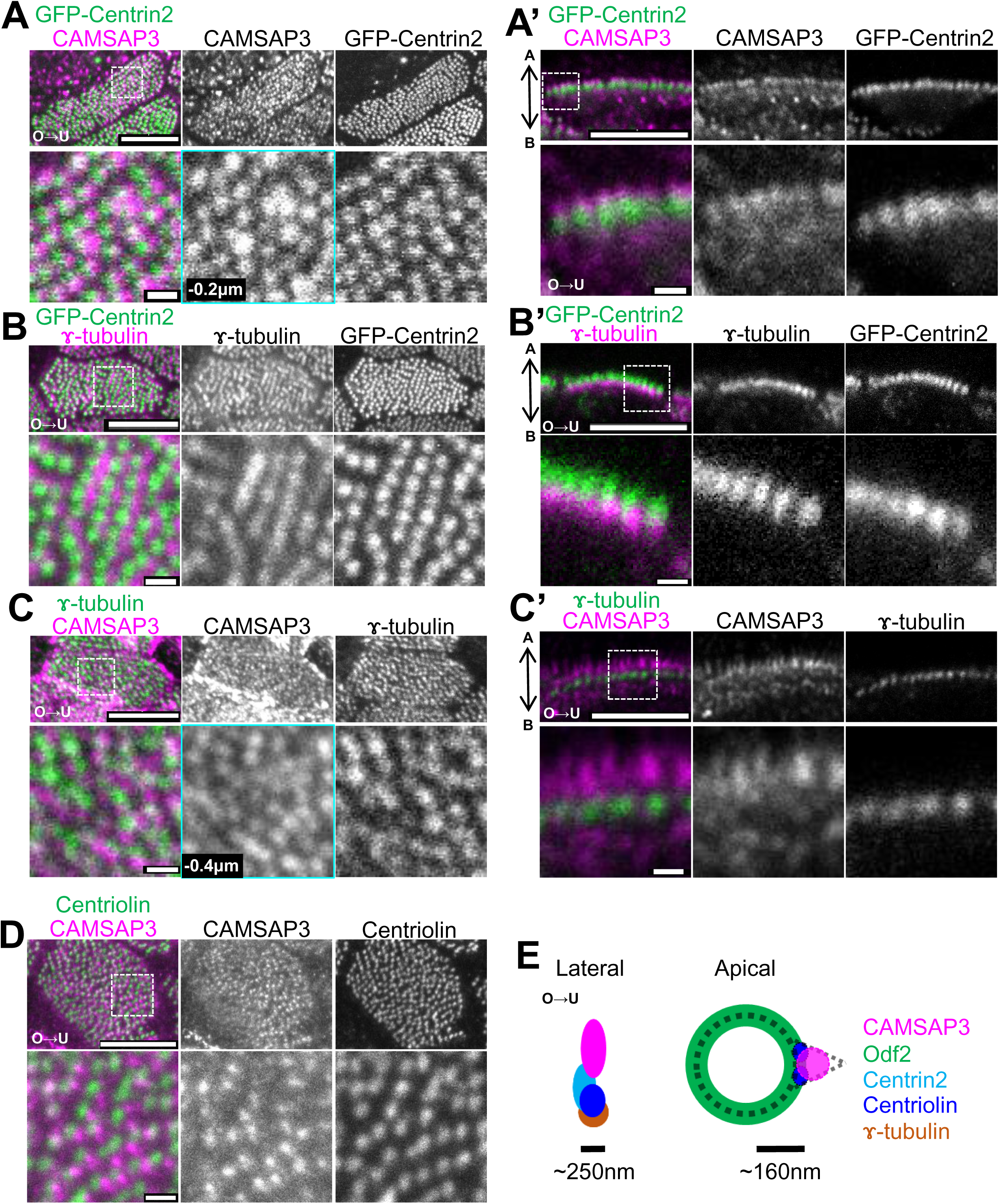
Localization of CAMSAP3 and other BB proteins in oviduct MCCs. Images taken from the apical side (A-D) and lateral side (A’-C’) of MCCs are shown. A and A’, Confocal images of the MCCs of adult *GFP-CETN2* (green) transgenic mouse stained with anti-CAMSAP3 antibody (magenta). A, Top, MIP image rendered from two images acquired at a step size of 0.2 μm. Bottom, magnified single plane images of the boxed region in A and A’. The focal plane of CAMSAP3 was 0.2 μm apical side to the plane of GFP-Centrin2, and is indicated as “-0.2 μm” (cyan frame). B and B’, Staining with anti–γ-tubulin antibody (magenta). B, Top, MIP image rendered from eight images acquired at a step size of 0.2μm. Bottom, Magnified images of the boxed region in B. C and C’, WT adult MCCs stained with anti-CAMSAP3 (magenta) and anti–γ-tubulin (green) antibodies. C, Top, MIP image rendered from six images acquired at a step size of 0.2 μm. Bottom, the focal plane of CAMSAP3 was 0.4 μm apical to the focal plane of γ-tubulin, and is indicated as “-0.4 μm” (cyan frame). D, Anti-Centriolin (green) and anti-CAMSAP3 (magenta) antibody staining. CAMSAP3 and Centriolin signals were acquired by STED and confocal microscopy, respectively. MIP image rendered from 13 images acquired at a step size of 0.2 μm. (Bar = 5 μm (upper panels) and 0.5 μm (lower panels), respectively (A-D and A’-C’)) E, Schematic diagram showing the localization of CAMSAP3 and other BB/BF proteins. Also see Figure S2.

The relative positions of CAMSAP3 and other BB and BF markers in mature MCCs are summarized in Fig. 3E. We observed that CAMSAP3 was located in the same direction as BF from the apical view, but was located apically when compared to other BF markers such as Centriolin and γ-tubulin.

### CAMSAP3 protein localizes to the BB prior to coordination of BB orientation

We went on to determine CAMSAP3 localization and the coordination of BB orientation during the maturation of MCCs. BB orientation has been reported to be coordinated in each cell at the final process when BBs spread over the entire apical surface during the maturation of MCCs in mouse trachea (Herawati et al., 2016). We began by observing the maturation process of MCCs of mouse oviducts (Fig. 4 and S3). The ratio of MCCs increased six weeks after birth and 80% of epithelial cells in the infundibulum of adult mouse oviduct were MCCs (Fig. S3A, A’) (Shi et al., 2014). We classified epithelial cells into five types based on the number and distribution of γ-tubulin and GFP-Centrin2 signals which are present in BBs. Similar progression patterns were observed when GFP-Centrin2 signals and anti–γ-tubulin staining signals were used to visualize BBs (Fig. S3B). Strong signals were observed for γ-tubulin (one or two large foci) and GFP-Centrin2 (multiple large foci) were observed around the apical surface in type I cells (Fig. 4A and Fig. S3B). Ring-shaped signals and condensed small dots were observed on the apical side of the nucleus in type II and type III cells, respectively. Unevenly scattered dots that formed clusters were observed around the apical surface of type IV cells, while dots were scattered across the apical surface in type V cells. Nuclear position and apical surface shape varied between cell types as indicated in Fig. 4A. In postnatal day 5 (P5) oviducts, we found that about 80% of cells were type I, and a few type V cells were also observed (Fig. S3A and A’). The ratio of type V cells increased as development progressed. Type IV cells were evident on P5, and the ratio of this cell type decreased after more than 2 weeks after birth. At P13, all cell types were observed. These analyses along the developmental stages suggested the progression of cell types from type I through type V. Time-lapse observation of *GFP-CETN2* mice also suggested the cell type transition from type I through type V (Fig. S3C and Sup. Mov.2). Additionally, we observed the appearance of multiple axonemes on the apical surface in type IV cells (Fig. S3D).

**Figure 4.**
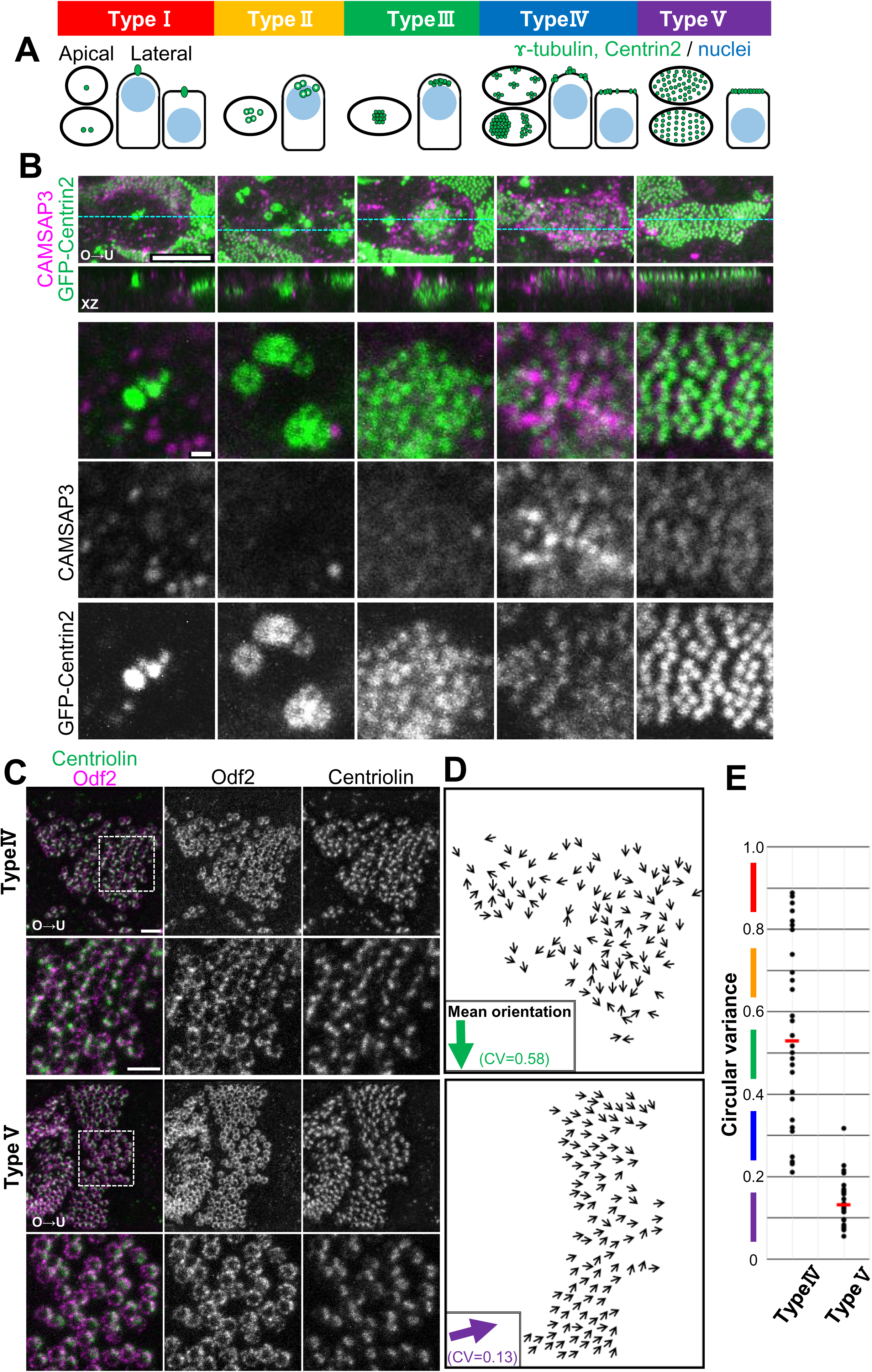
CAMSAP3 protein localization during maturation of MCCs. A, Schematic diagram of cell types observed on P13. Cells were classified into five types based on cell shape, nuclear position, and GFP-Centrin2 or γ-tubulin distribution. B, Confocal images of GFP-Centrin2 (green) and anti-CAMSAP3 antibody staining (magenta). Images were acquired and processed under the same conditions to enable direct comparison of signal intensities. Top, MIP images rendered from 20 images acquired at a step size of 0.2 μm. Panels in the second row show xz images along the blue dotted lines in the top row. Bottom three panels show the single focal plane where the GFP-Centrin2 signal was observed. See also Figure S3. C, Super-resolution microscopic images of P13 WT cells stained with anti-Centriolin (green) and anti-Odf2 (magenta) antibodies. MIP images rendered from 15 images acquired at a step size of μm after each image was subjected to displacement correction (Fiji menu command Correct 3D Drift) (Bar = 1 μm). Magnified images of boxed regions in the top panels are shown on the bottom (Bar = 0.5 μm). D, The direction of BB orientation. Number of BB analyzed and CV in the representative images: n = 108 and CV = 0.58 (type IV); n = 107 and CV = 0.13 (type V). E, Distribution of CVs in type IV and type V cells. Each dot in the graph corresponds to one cell, and the red bar represents the median. Three animals were analyzed. Type IV: 28 cells (average number of BB analyzed/cell = 97; median CV = 0.53 indicated with a red bar). Type V: 28 cells (average number of BB analyzed/cell = 116; median CV = 0.13).

Next, we analyzed when CAMSAP3 became co-localized with the BB components during the development of MCCs. In P13 oviducts, CAMSAP3 showed dot-like signals around apical surface in all cell types (Fig. 4B). In type IV and type V cells, CAMSAP3 was observed adjacent to GFP-Centrin2. CAMSAP3 signal intensity was stronger in type IV than in type V cells, and CAMSAP3 dots were widely distributed over the apical surface while GFP-Centrin2 formed clusters of dots. In type V cells, CAMSAP3 localized at BBs, which was similar to their observed localization in adult MCCs, while CAMSAP3 dots were even smaller when compared to those in adult MCCs. These results suggest that CAMSAP3 localizes to BBs both during and after BB docking to the cell surface.

We also examined the localization of Odf2 and Centriolin using P13 mouse oviducts (Fig. S3E and F). Centriolin colocalized with GFP-Centrin2 in all cell types; however, Odf2 and Centriolin colocalized only in type IV and type V cells (Fig. S3F). Odf2 localized to BBs in 83% of type IV cells, and 100% of type V cells. The above results indicate that CAMSAP3 localizes to the BB at a later time than other BB/BF components, including Centrin2, Centriolin, γ-tubulin, which were already colocalized before BB docking. Odf2, however, localizes to BBs later than CAMSAP3.

We next focused on type IV and type V cells and determined BB orientation (Fig. 4C-E) as described above. In type IV cells, individual BB were oriented in various directions within each cell, and CV values varied widely between 0.2 and 0.9 (n = 28 cells, median = 0.53). In type V cells, however, BBs were oriented in a similar direction, and the CV value of most cells was smaller than 0.32 (n = 28 cells, median = 0.13) (Fig. 4E). Thus, BB orientation was aligned during the transition from type IV to type V. Taken together, these observations revealed that CAMSAP3 localizes to the BBs during their docking to the plasma membranes prior to the coordination of BB orientation.

### BB orientation is not coordinated in *Camsap3 dc/dc* mutant

To determine whether CAMSAP3 plays a role in coordination of BB orientation, we examined MCCs in the *Camsap3 dc/dc* mutant (Toya et al., 2016) oviduct. We found that multicilia formed in *Camsap3 dc/dc* on the luminal surface of the oviduct epithelium (average = 221 BBs/cell, SD = 33.3, n = 20 cells, two animals). The distribution of BBs in each mutant MCC was not aligned compared to WT (Fig. 5A and A’). The BB orientation in each cell was not in the same direction in the mutants (Fig. 5B and B’), and CV values varied from 0.1 to 0.9. Compared to the CV in WT litter mates (median = 0.16, n = 10 cells, average = 195 BBs/cell analyzed), the CV values were significantly higher in *Camsap3 dc/dc* (median = 0.47, n = 20 cells, average 182 BBs/cell analyzed; Fig. 5C and S5). The CV value for *Camsap3 dc/dc* was also significantly higher than that of the *Celsr1-/-* mutant and comparable to WT immature type IV cells (Fig. S5). BBs were distributed across the entire apical surface in *Camsap3 dc/dc*, which is different from WT immature type IV MCCs, whereas BBs formed clusters, suggesting that high CV is not due to the delay in MCC maturation. Then, we compared the mean BB orientation between adjacent cells against the longitudinal axis of the oviduct, finding that the mean BB orientation of each cell was aligned between cells, even between cells with higher CV values (Fig. 5 D and D’). The mean BB orientation was also aligned with the direction of the oviduct folds (data not shown).

**Figure 5.**
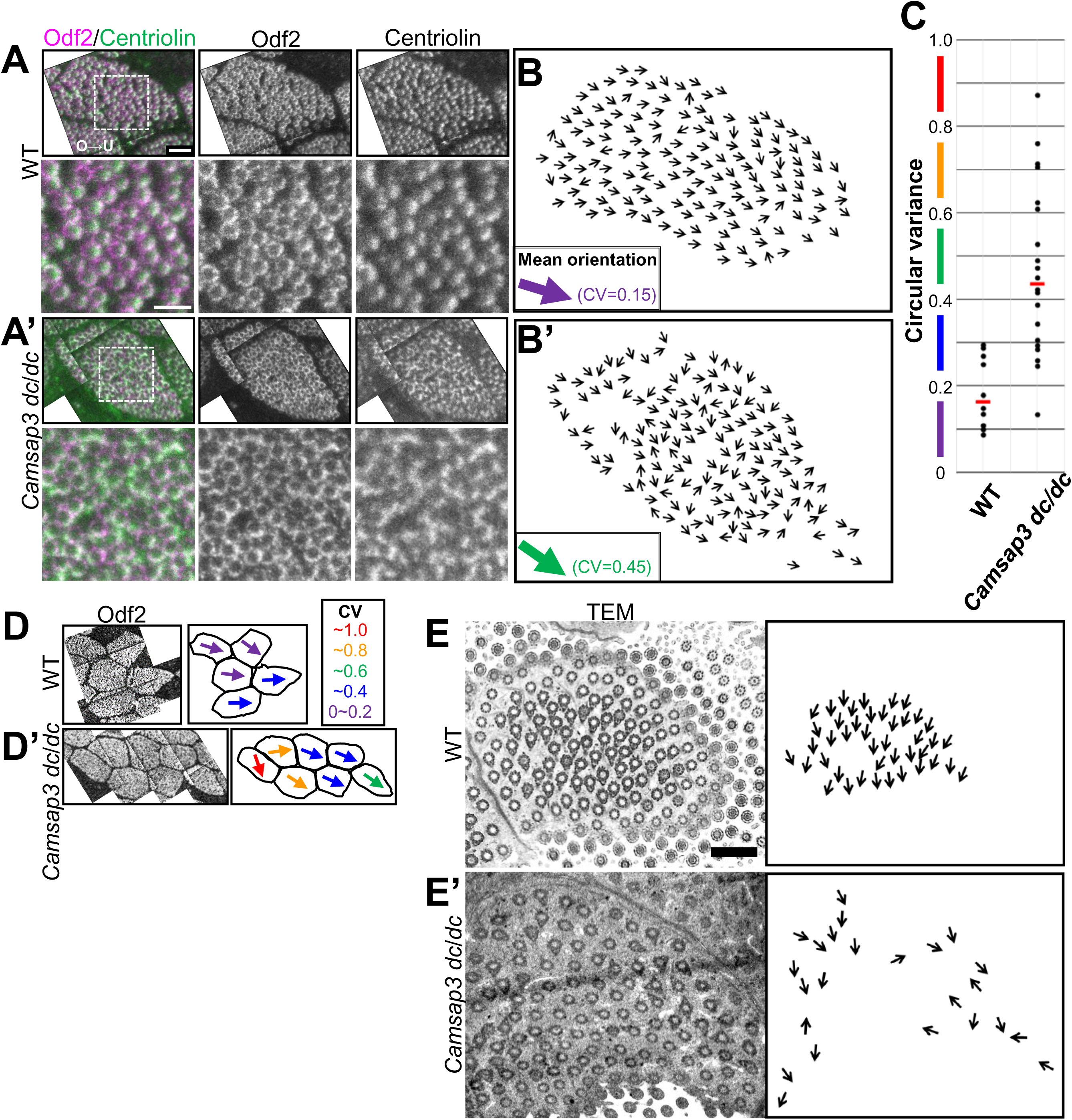
BB orientation is not coordinated within each *Camsap3 dc/dc* cell. A and A’, Super-resolution microscopy (STED) images of adult WT (A) and *Camsap3 dc/dc* mutant (A’) MCCs stained with anti-Odf2 (magenta) and anti-Centriolin (green) antibodies. MIP images rendered from 10-15 images acquired at a step size of 0.1 μm; (Bar = 1 μm (upper panels) and 0.5 μm (lower panels)). B and B’, Orientation of BBs in A and A’. In these images, 148 BBs were analyzed and CV = 0.15 in WT, and 148 BBs and CV= 0.45 in *Camsap3 dc/dc*. C, CV of BB orientation in each cell. Each dot in the graph corresponds to one cell, and the red bar represents the median. Ten WT cells (average number of analyzed BBs/cell = 195) and 20 *Camsap3 dc/dc* cells (182 BBs/cell) were analyzed. The datasets are identical to those in Figure S5. D and D’, Comparison of mean BB orientation in WT (D) and *Camsap3 dc/dc* (D’). The direction of the mean BB orientation of each cell is shown with arrows, and color corresponds to the CV value of each cell. E and E’, TEM images of BB and BF in WT and *Camsap3 dc/dc*. In the right panels, arrows indicate BB directions traced manually from the EM images (Bar = 1 μm).

Electron microscopy observations also revealed BB orientation defects in the *Camsap3 dc/dc* mutant (Fig. 5E and E’), and were consistent with the results obtained using super-resolution microscopy. In a minor population, we observed structures similar to BFs protruding from multiple positions on BBs (Fig. S4A). These results indicate that CAMSAP3 is indispensable for aligning BB orientation in each cell.

### Planar polarized protein localization is maintained in *Camsap3 dc/dc*

Next, we examined if PCP is maintained in *Camsap3 dc/dc* mutant oviducts (Fig. 6A and A’). Cell shape was elongated and epithelial folds formed similarly to WT. Celsr1 and Vangl1 localized to the cell boundary perpendicular to the ovary-uterine axis in a polarized fashion in the same manner as in WT when they were quantified (Fig. 6B and B’). The enrichment of polarized microtubules near the cell boundary on the uterine side was also evident (Fig. 6C and D), as seen in WT mice (Fig. 2A). Thus, despite the disruption of the coordinated BB orientation, PCP factors and microtubule enrichment occurred in a normal orientation in the mutant oviducts. These results suggest that the *Camsap3 dc/dc* mutant phenotype is not due to the loss of PCP at the tissue level.

**Figure 6.**
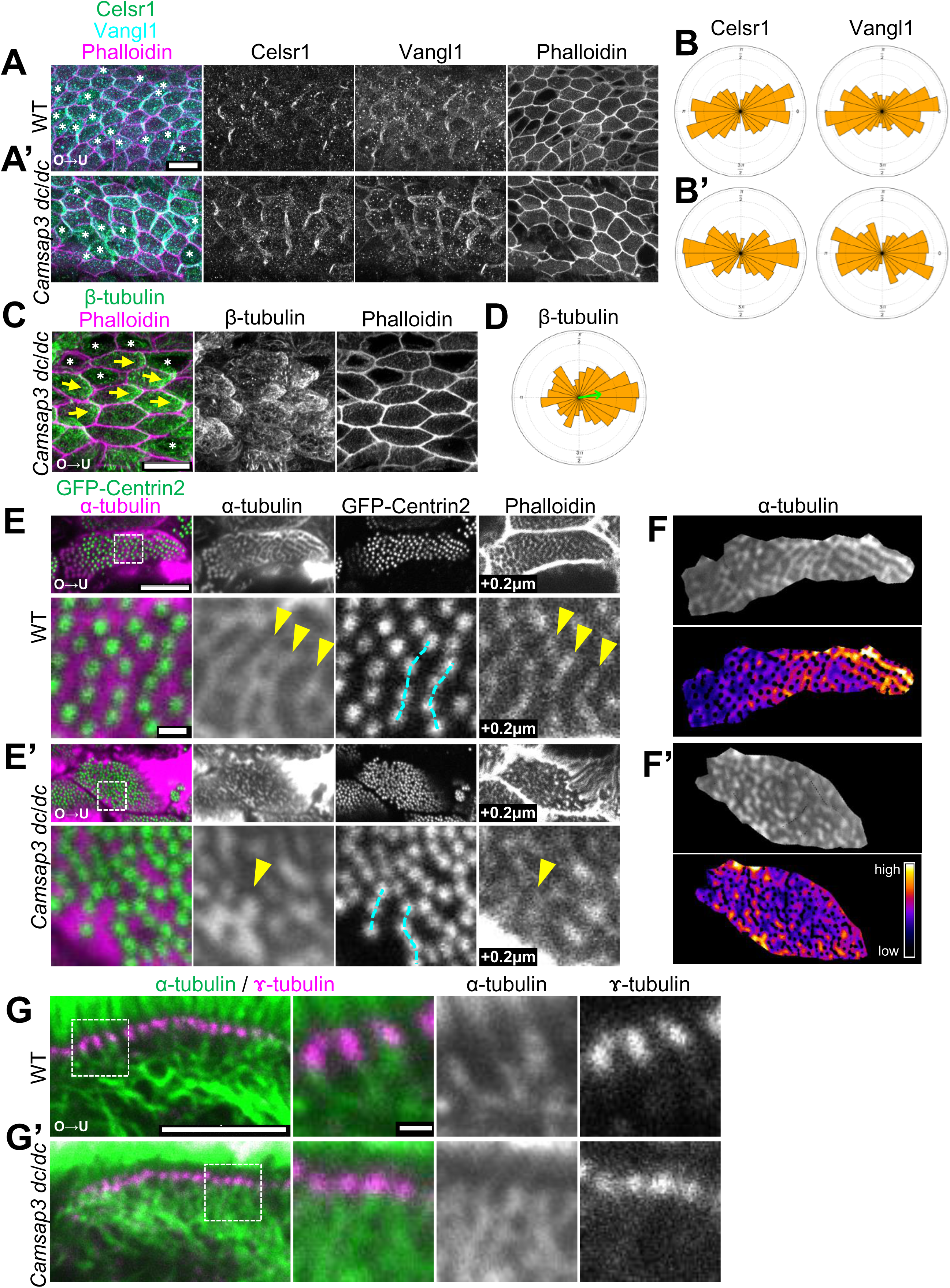
Localization of proteins that regulate PCP is similar to wild type but the distribution of microtubules around BBs was altered in *Camsap3 dc/dc* mutant. A and A’, Confocal microscopy images of adult oviducts of littermate control (WT) and *Camsap3 dc/dc* mutant stained with anti-Celsr1 (green), anti-Vangl1 (cyan), and phalloidin (magenta). MIP image rendered from 15 images acquired at a step size of 0.2 μm (Bar = 10 μm). B and B’, Rose diagram showing the results of quantifying Celsr1 and Vangl1 localization. The angle was classified into 24 bins. The area of each bin is proportional to the number of cells distributed in each bin. A total of 99 WT cells and 119 *Camsap3 dc/dc* cells were analyzed. Secretory cells are indicated with asterisks. C, Confocal images adult of *Camsap3 dc/dc* oviduct stained with anti–β-tubulin antibody (green) and phalloidin (magenta). MIP images rendered from 20 images acquired at a step size of 0.2 μm. The directions of β-tubulin enrichment are shown with yellow arrows. Secretory cells are indicated with asterisks (Bar = 10 μm). D, Rose diagram showing the quantified direction of β-tubulin signal enrichment. E and E’, Apical views of the oviducts from littermate WT (E) and *Camsap3 dc/dc* (E’) mice showing GFP-Centrin2 against background stained with anti–α-tubulin antibody and phalloidin. Magnified images of boxed regions in the upper panels are shown at the bottom. Images of two representative focal planes are shown. The focal plane of phalloidin signals were 0.1 μm basal (shown as “+0.2 μm”) to the focal plane of α-tubulin and GFP-Centrin2 signals. Note that striped microtubule signals are located to the side of aligned BBs (highlighted by cyan dotted lines) in WT but not in *Camsap3 dc/dc*. Distribution of microtubules and phalloidin signals are comparable in WT but not in *Camsap3 dc/dc* (compare signals pointed by yellow arrowheads); (Bar = 5 μm (upper panels) and = 0.5 μm (lower panels)). F and F’, Reconstructed images of α-tubulin signals located at the level of GFP-Centrin2 signals. Different focal planes were merged for the reconstruction because the apical surface was not flat. Intensity of signals were color-coded as shown in the bottom right inset. Position of GFP-Centrin2 signals were masked to highlight the distribution of microtubules between BBs. G and G’, Lateral views of WT (G) and *Camsap3 dc/dc* (G’) MCCs stained with anti–α-tubulin (green) and anti–γ-tubulin (magenta) antibodies; (Bar = 5 μm), Magnified images of boxed region in the left panels are shown on the right; (Bar = 0.5 μm).

### Apical microtubule distribution is disturbed in *Camsap3 dc/dc* mutant MCCs

To elucidate the mechanism of how CAMSAP3 coordinates cilia orientation within cells, we examined the distribution of microtubules more closely in *Camsap3 dc/dc* mutant MCCs, as it might be related to the disorganization of BB orientation. In *Camsap3 dc/dc,* α-tubulin was distributed to fill the spaces between GFP-Centrin2 signals as observed in WT. Notably, however, microtubule stripes became fragmented in the mutant cells (Fig. 6E’), although some of BBs were still aligned into rows (Fig. 6E and E’ dotted lines). The analysis of the overall staining intensity of α-tubulin located at the level of GFP-Centrin2 in individual cells (in the range of 0.6 μm) showed that, while α-tubulin was enriched at the uterine side of WT cells, this enrichment was no longer observed in *Camsap3 dc/dc* (compare α-tubulin panels in Fig. 6F and F’), although other microtubule populations still exhibited a gradient distribution as shown in Fig. 6C. We next examined the microtubules, which extend apico-basally into the cytoplasm from BBs, from the lateral side (Fig. 6G and G’), finding that the assembly of this population of microtubules looked normal in *Camsap3 dc/dc* mice. We also checked for acetylation of microtubules by staining with anti-acetylated tubulin antibody, but we did not detect any differences between WT and mutant mice (Fig. S4D and D’). These observations suggest that the assembly of microtubules horizontally associated with BBs are selectively regulated by CAMSAP3.

As a previous study showed that CAMSAP3 binds actin filaments via ACF7 (Ning et al., 2016), we observed actin distribution using phalloidin. Phalloidin signals also formed stripes about 0.2 μm basal from the plane of α-tubulin stripes in WT (Fig. 6E), whereas, in *Camsap3 dc/dc*, the distributions of α-tubulin and phalloidin signals became less identical (Fig. 6E’ arrowheads). At the apical surface of MCCs, phalloidin-positive protrusions were observed more frequently in the mutant cells (Fig. S4B and B’). A small number of these protrusions showed a cyst-like structure, and the cysts occasionally contained microtubes or multiple axonemes (Fig. S4D’ and C), which are never seen in WT cells. These suggests that actin filament organization was also affected as a result of *CAMSAP3* mutation. Finally, we observed the distribution of CAMSAP3 in *Celsr1*-/- MCCs, and found no change in its distribution after *Celsr1* loss (Fig. S6A and A’), implying that the Celsr1 and CAMSAP3 systems differentially regulate BB orientation.

## Discussion

The present study analyzed BB orientation and its coordination between cells in the oviducts using two mutants which have lost Celsr1 or CAMSAP3 functions. Each mutant displayed distinct abnormalities concerning these processes. In *Celsr1* mutants, PCP was partly abrogated, whereas the intracellular polarity represented by BB orientation and microtubule gradient were normally maintained within individual cells. On the other hand, *CAMSAP3*-mutated cells showed disruption of BB orientation in individual cells, but not losing PCP of the cell sheet. These observations suggest that BB orientation in individual cells occurs independently of Celsr1, and cooperation of the Celsr1 and CAMSAP3 systems is necessary to determine the entire polarity of oviducts in both cellular and tissue levels.

### Ciliary movements are directly linked to the physiological activities of organs they are belong to

Fluid flow in the oviduct/fallopian tube was recently reported to be directed from the uterine side to the ovarian side by the secretion of fluid and contractility of smooth muscles in the isthmus of the oviduct (Hino and Yanagimachi, 2019). Eggs are transported steadily against the direction of fluid flow from the infundibulum to the ampulla by directional movements of the cilia, which are coordinated both within individual cells and between cells. The flow of cerebrospinal fluid (CSF) in the ventricles also depends largely on the movement of cilia belonging to ependymal cells, but flow in the ventricles also shows a complicated pattern (Eichele et al., 2020; Faubel et al., 2016). Additionally, the direction of cilia is not always the same as the direction of the organ. In this study, observing BBs with super-resolution microscopy allowed us to determine whether cilia orientation is coordinated between cells even in the wide field within the folds of the oviduct epithelium. As the folds are along the tube in the infundibulum, cilia orientation is aligned with organ orientation, i.e. the longitudinal axis of the oviduct.

### Celsr1 coordinates cilia orientation between cells in the oviduct

The mechanism for coordinating the orientation of multicilia differs between animal species and between organs. Boutin and colleagues have reported that the functions of PCP factors also vary (Boutin et al., 2014). In the mammalian ependyma, one group of PCP factors (Celsr2, 3, Fzd3, and Vangl2) regulates cilia orientation within individual cells while another group (Celsr1, Fzd3, and Vangl2) regulates cilia orientation between cells (tissue polarity). Either microtubule enrichment or anchoring to the cell boundary was also observed in ependymal cells, and the direction of enrichment was less coordinated between cells in *Celsr1-/-*. The involvement of Celsr1 in the coordination of cilia orientation may differ between ependyma and oviduct epithelium. While cilia patch displacement and microtubule enrichment were correlated in *Celsr1-/-* ependyma, the orientation of patch displacement did not correlate with mean cilia orientation. In contrast, the direction of microtubule enrichment was not coordinated between oviduct epithelial cells in *Celsr1-/-* while the mean BB orientation coincided with the direction of microtubule enrichment. Among the Celsr family proteins, Celsr1 probably plays a major role in the oviduct epithelium because Celsr3 expression was not detected in oviduct epithelial cells, and *Celsr2* mutants have no obvious ciliary phenotype in the oviduct. Cilia orientation was not completely random in individual cells lacking Celsr1, but the orientation between cells was not the same. This difference in cilia orientation suggests that Celsr1 is mainly responsible for coordinating cilia orientation between cells.

### Intracellular self-organizing mechanisms that coordinate cilia orientation in individual cells

Cells that lack Celsr1 have been proposed to retain the ability to self-organize the distribution and orientation of cilia within individual cells. But how is cilia orientation coordinated within individual cells? Cytoskeletal elements are known to play central roles in regulating cilia orientation in cells. The type of cytoskeletal element involved in such regulation depends on the animal species and the specific organ (Sandoz et al., 1988; Werner et al., 2011; Clare et al., 2014; Antoniades et al., 2014; Chevalier et al., 2015; Herawati et al., 2016; Tateishi et al., 2017;, Vu et al. 2019). In mammalian MCCs found in the ependyma and trachea, microtubules are considered important for aligning cilia orientation within the cells. microtubules that localize just below the apical surface are connected to BFs. Microtubules are also enriched on one side of cells in a region that is slightly deeper than the apical microtubule meshwork around the cell-cell boundaries. We found that this is also common in oviduct MCCs. Moreover, we observed that microtubules are concentrated on the uterine side of each cell, and the apical microtubule meshwork forms a striped pattern with a gradient from the uterine side along the longitudinal axis of the oviduct. We found that in the *Celsr1-/-* mutant, microtubule stripes were present between the BBs, and microtubule enrichment was also conserved in more than 75% of MCCs. BBs were oriented toward the site of microtubule enrichment in these cells. These results suggest that Celsr1 is not essential for intracellular microtubule enrichment and apical stripe formation. We cannot rule out the possibility that other PCP factors may coordinate cilia orientation within each cell. However, even when *Vangl2* is deficient, ciliated cells do not show visible abnormalities (Usami, unpublished data), in contrast to ependymal cells. Therefore, MCCs of the oviduct may be less dependent on PCP factors.

### Microtubule networks and the regulation of cilia orientation by CAMSAP3

We found that CAMSAP3, a non-centrosomal microtubule minus-end binding protein, localizes to the base of the cilia of MCCs, in addition to γ-tubulin, a centrosomal microtubule minus-end binding protein (Clare et al., 2014). The localization of CAMSAP3 to BB of the trachea was also recently reported (Robinson et al., 2020). In contrast to the prevailing view that a connection of microtubules to a BF controls cilia orientation, we found that CAMSAP3 was not localized to BF. Instead, its localization is more apical and contributes to the alignment of cilia orientation. γ-tubulin has been shown to be localized in BF, and we found that CAMSAP3 also exists in the same direction as γ-tubulin in each BB in the plane near the apical surface. The apical microtubule stripes and CAMSAP3 are on the same plane, and CAMSAP3 may interact with microtubules in this plane. During the maturation of MCCs, γ-tubulin exists in the BB regardless of the developmental stage, but CAMSAP3 localizes to BBs at the stage of BB docking. Whereas the subcellular localization of γ-tubulin, Centrin2, and Centriolin were similar at all stages, CAMSAP3 was present on the apical region throughout all stages. CAMSAP3 only localizes to BBs after BB is docking. From the above, it is suggested that γ-tubulin and CAMSAP3 may play different roles in organizing microtubules in MCCs. In the intestinal epithelium, the absence of CAMSAP3 affected microtubules running from basal and attach apical surface (Toya and Takeichi, 2016), whereas in the oviduct, the microtubules running apicobasally were maintained in *Camsap3 dc/dc* mutant MCCs. On the other hand, the distribution of apical microtubules around BB was affected. We speculate that the minus-end of microtubules running apicobasally may be bound to γ-tubulin in MCCs, but this needs to be elucidated in the future.

Odf2 has been suggested to localize to BBs late in MCC maturation following to CAMSAP3 localization to the BBs. Both CAMSAP3 and Odf2 may become localized to BBs when the polarity or asymmetry of BBs is established. In *Odf2* mutant tracheal MCCs, the BF structure is also lost, and Odf2 is essential for the polar formation of BB (Kunimoto et al., 2012). CAMSAP3 is, however, dispensable for BF formation, because the BF structure can still form.

### Why is cilia orientation disturbed in the *Camsap3* mutant?

It is unlikely that the cells lacking CAMSAP3 could not acquire planar polarity except for cilia orientation, because the PCP factor localizes correctly, microtubules are enriched on the cell edge, and folds are formed along the longitudinal axis of the oviduct. BB/BF components (Odf2, Centriolin, Centrin2, γ-tubulin) localize to BBs. The formation of BBs and BFs is almost normal except for some minor populations. We observed some BBs with multiple BFs, and CAMSAP3 may also be responsible for limiting BF to one.

The BB orientation defect is quite likely due to local abnormalities around BBs. To align BBs and coordinate orientation of BBs, BFs should be connected in the same orientation by cytoskeletal elements. One possibility is that CAMSAP3 is involved in anchoring BFs to microtubule networks. We observed the impaired accumulation of microtubules in the apical region around BBs in *Camsap3 dc/dc* mutant MCCs. We speculate that stable apical microtubule stripe formation is disrupted or the stability of apical microtubules are defective in the absence of CAMSAP3. Due to this change, BFs cannot anchor stably to the cytoskeleton, and the orientation of BBs are not aligned. The binding of CAMSAP3 and ACF7 may mediate the interaction between microtubules and actin filaments (Ning et al., 2016; Noordstra et al., 2016) and this interaction may also help microtubules to accumulate around BBs and connect neighboring BFs in the same orientation. The stable accumulation of apical microtubules connecting BFs may therefore be the major function of CAMSAP3.

### The mechanism for aligning cilia orientation within and between cells

In this study, we propose that the orientation of multicilia is regulated at multiple scales. The direction of cilia has been suggested to be autonomously aligned within individual cells by either controlling the binding of BFs to microtubules or by regulating the distribution of stable apical microtubules, regulated by CAMSAP3. Celsr1 is a PCP factor that aligns microtubule networks between cells to coordinate tissue-wide cilia orientation. We tried to analyze interaction between these two mechanisms by making double knockout, but we could not obtain the *Camsap3 dc/dc*; *Celsr1 -/-* mice because they were perinatal lethal. In addition to these two hierarchies, microtubules may be enriched at cell boundaries in a polarized fashion, and therefore form a concentration gradient inside cells that may connect these two systems. How these multi-scale mechanisms are connected so that the orientation of cilia in the whole tissue is aligned should be elucidated in the future.

## Acknowledgments

This work was supported by JSPS KAKENHI Grant Number 25291054, 15H01220, 17H03689, 16H06280, and 19K16153. T.F. was supported by JST CREST, JPMJCR1654. We thank Holden Higginbotham and Hiroshi Hamada for the *GFP-Centrin2* mouse; Sachiko Tsukita for the anti-Odf2 antibody and hybridoma; Masahiko Itoh and Mikio Furuse for the anti-ZO1 antibody; Yoshikatsu Sato and Tetsuya Higashiyama (ABiS) for technical support and advice concerning STED microscopy, and Gen Yamada for TEM support; Hiroko Saito, Azusa Kato, Tetsuhisa Otani, and other lab members for technical support with animal care and helpful discussions. We also thank ExCELLS for allowing us to use STED. This study was also supported by the Spectrography and Bioimaging Facility (NIBB) and the EM facility (NIPS).

## Author Contributions

Conceptualization, F.M.U., M.A., and T.F.; Methodology, F.U., M. A., D.S., and T.F.; Software, M.A. and D.S.; Investigation, F.M.U., D.S and Y.H.; Resources, S.O., Y.H., F.T., and M.T.; Writing, F.M.U., M.A., M.T, and T.F.; Visualization, F.M.U. and M.A.; Funding Acquisition, M.A. and T.F.; Supervision, M.T. and T.F.; Project Administration, T.F.

## STAR Methods

### Animals

Female mice (Slc; ICR, Japan SLC) were used to study MCCs in WT. *Celsr1-/-* mutant mice (Ravni et al., 2009), *Camsap3 dc/dc* mutant mice (Toya et al., 2016), and *GFP-Centrin2* (*GFP-CETN2*) Tg mice (Higginbotham et al., 2004) were described previously. Animal care and experiments were conducted in accordance with the Guidelines of Animal Experimentation of National Institutes for Natural Sciences and RIKEN animal experimentation guidelines. All animal experiments were approved by either the Animal Research Committee of National Institutes for Natural Sciences or by the Institutional Animal Care and Use Committee of Riken Kobe Branch respectively. Animals were maintained in a light- and temperature-controlled room using a 12 h light:12 h dark cycle at 23+/-2 °C.

### Immunohistochemistry

The fluorescent images were taken by using a Nikon A1 confocal microscope (Nikon) and TCS SP8 STED (Leica). Sample preparations were basically the same as previously reported (Shi et al., 2014). Anti-Odf2 antibody–producing hybridoma was a gift from S. Tsukita (Tateishi et al., 2013). All primary antibodies and secondary antibodies used in this study are listed with fixation conditions in key resource table.

After 4% PFA fixation, samples were treated with 0.1% TritonX-100 in PBS at room temperature (RT) for 1 h. Fixed samples were blocked with Blocking One (Nacalai Tesque) at RT for 1 h, and incubated with primary antibodies in Blocking One at 4 °C overnight. Incubation with α- or β-tubulin antibodies was for 36 h at 4 °C. After washes with PBS, samples were incubated with secondary antibodies at RT for 3 h or 4 °C overnight, or 24 h at 4 °C for α- or β-tubulin antibodies staining. After washes, each sample was mounted in Fluoromount-G (SouthernBiotech). ProLong Diamond (Thermo Fisher Scientific) was used to mount samples for STED observation. The ovarian side of the oviduct was determined by the presence of fimbriae.

### Quantifying the localization of PCP proteins and microtubule enrichment

Localization of PCP proteins was quantified by calculating nematic order as described (Shi et al., 2014; Aigouy et al., 2010). Maximum intensity projection (MIP) images were rendered from five serial images acquired with a step size of 1 μm. The orientation of microtubule enrichment was determined by using this method with minor modifications. Since microtubules were enriched at the proximity of cell boundaries, cell boundaries were labeled with phalloidin and microtubule signals within the range of 0 to 5 pixels away from the cell boundaries were extracted using ImageJ. Nematic order defines only the axis and the magnitude of polarity. Additionally, the vector of microtubule polarity was determined by separating microtubule signals into two halves with a line perpendicular to the axis of the polarity, and by comparing intensities between these two separated microtubule signals. MIP images were rendered from twenty serial images acquired with a step size of 0.2 μm.

### Quantifying basal body orientation

BB orientation was determined by Odf2 and Centriolin signals after acquiring images with TCS SP8 STED (Leica). All images were acquired at a step size of 0.1 μm, and images from the Odf2 channel were subjected to an ImageJ macro (Correct 3D Drift). Same parameter of “Correct 3D drift” was applied for images of Centriolin. On each Odf2 ring, an arrow pointing to the center of ellipse/two points of Centriolin signals (as shown in Fig. 1A’) was manually plotted. Making use of shape anisotropy of the arrow, the angle between arrow and the longitudinal axis (ovary > uterus) of the oviduct was measured by Shape Descriptor Map (BioVoxxel_Toolbox; Fiji). In each cell, the CV of each set of angles was calculated by using R as described previously (Zar, 2010; Shi et al., 2014). To acquire images using a transmission electron microscope (JEM-1010, Jeol; equipped with 2k x 2k CCD camera, Olympus Soft Imaging Solutions), sample blocks were prepared as described (Shi et al., 2014). Ultrathin sections (50–70 nm) were made with a microtome (Leica EM UC7), and images were obtained on a grid (OkenShoji150-D, 75-A mesh). To measure the orientation of BB in TEM images, an arrow was drawn manually from the center of BB to BF. CV was calculated as described above.

### Time-lapse recordings of MCC maturation

The oviducts of *GFP-CETN2* Tg mice (postnatal day 13) were opened longitudinally, and were put into a drop of Dulbecco’s modified eagle’s medium with high glucose (Gibco#31053) containing 10% FBS on a glass bottom dish (MatTek P35G-1.5-10-C). The drop was covered with liquid paraffin (128-04375, Wako) to avoid evaporation. The apical surface of luminal epithelium faced the bottom of the dish. Oviducts were cultured at 37 °C under 5% CO_2_ during the recording, and imaged with a spinning disc confocal system (Cell voyager CV1000; Yokogawa) using a 100x objective lens (UPLSPAPO100XS, Olympus) at 1 h intervals. At each timepoint, 40 slices were acquired at a step size of 0.5 μm.

## Supplemental Information

### Supplemental Figure legends

**Fig. S1.**
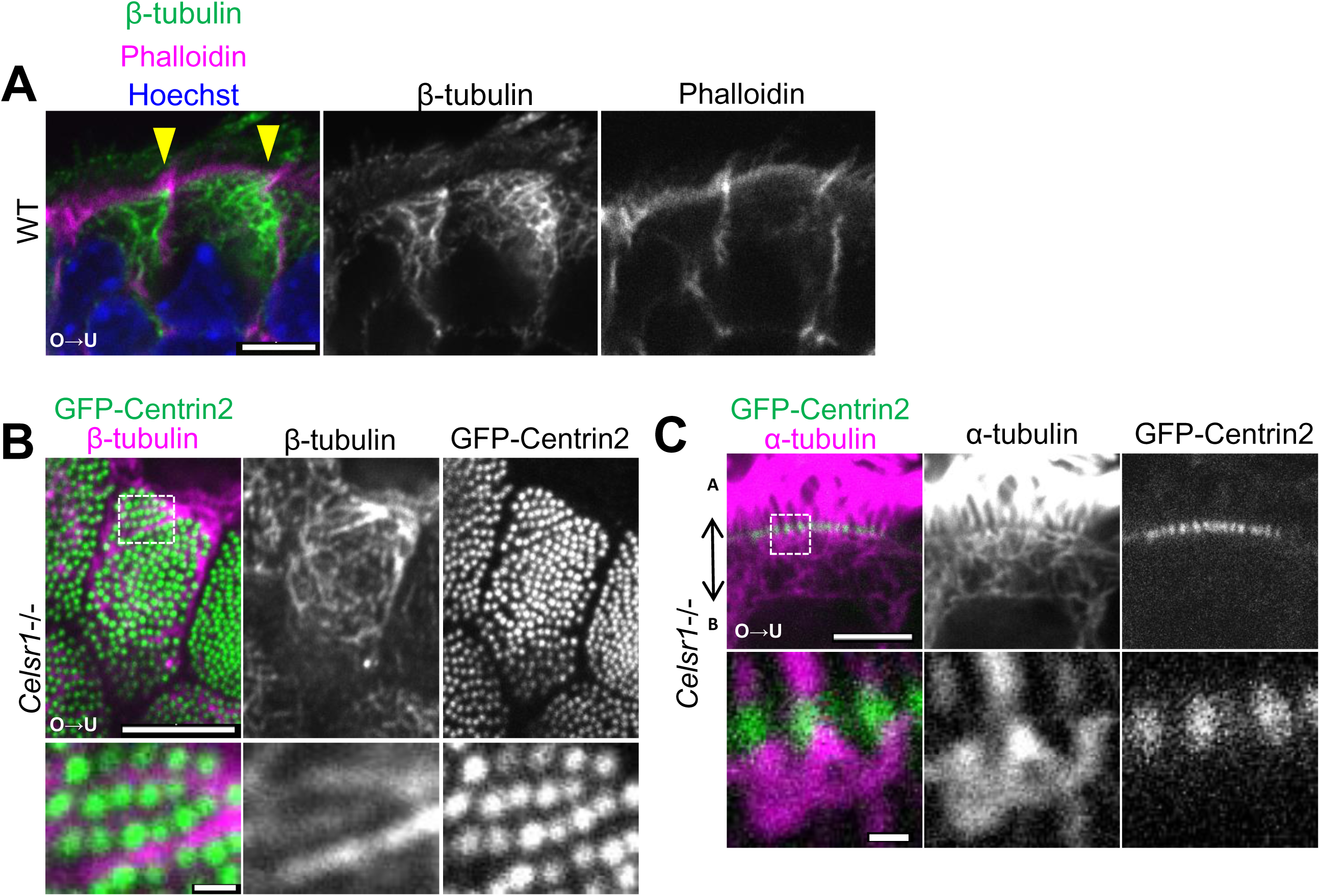
Microtubule organization was maintained in *Celsr1*-/- MCC. A, Lateral view of WT MCCs stained with anti–β-tubulin (green) and phalloidin (magenta). Arrowheads indicate cell-cell boundary identified by phalloidin signals. (Bar = 5 μm) B and C, Apical (B) and lateral view (C) of MCCs of *Celsr1-/- ; GFP-CETN2* transgenic mouse stained with anti–β-tubulin (magenta in B) or anti–α-tubulin antibodies (magenta in C). The ovarian and uterine side are on the left and right, respectively in all images (shown as O > U). A, Top row, MIP image was rendered from three images acquired at a step size of 0.2 μm. Bottom row shows high magnification images of a single plane. C, Single plane images. Bottom row, high magnification. Apical is to the top of the images. (Bar = 5 μm and 0.5 μm in the top and bottom rows, respectively, in both B and C.)

**Fig. S2.**
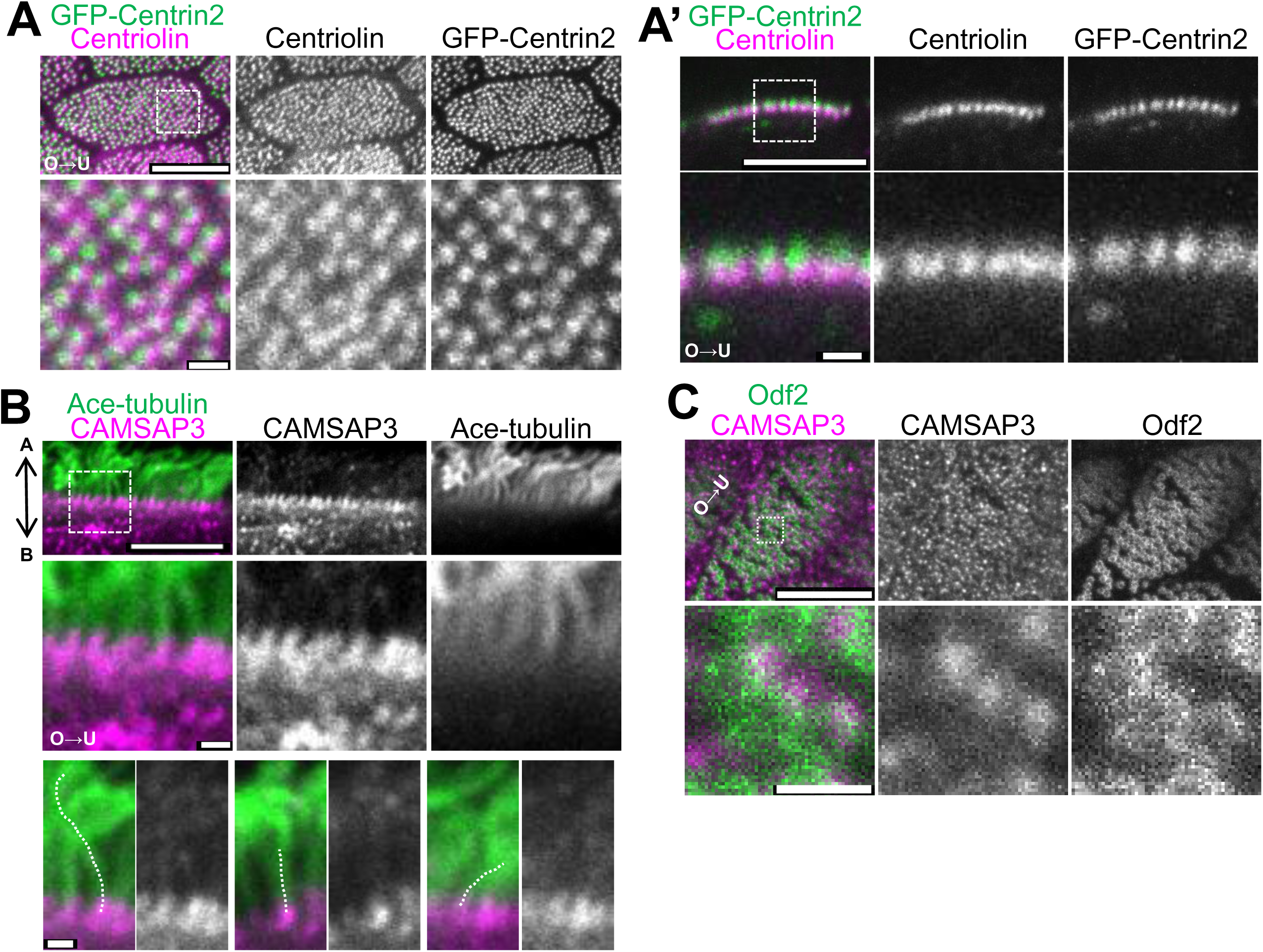
Localization of BB related proteins. A and A’, Apical (A) and lateral (A’) views of *GFP-CETN2* (green) tg oviduct stained with anti-Centriolin (magenta) antibody. MIP images rendered from 5 images acquired at a step size of 0.2 μm (A), and single plane images (A’). Lower panels show magnified views of the boxed region in the upper panel. (Bar = 5 μm (top), = 0.5 μm (bottom)) B, Lateral view of adult WT oviduct stained with anti-CAMSAP3 (magenta), and anti-acetylated tubulin (green) antibodies. Middle panels show magnified images of the boxed region in the upper panel. Bottom panels show cilia (dotted lines) standing with different angles from the apical surface. (Bar = 5 μm (top panels), 0.5 μm (middle panels), and 0.5 μm (bottom panels)) C, Super resolution images (obtained by STED) of WT oviduct stained with anti-CAMSAP3 (magenta) and anti-Odf2 (green) antibodies. MIP image was rendered from three images acquired at a step size of 0.2μm. Bottom panels show the magnified images. (Bar = 5 μm (top) and 0.5 μm (bottom))

**Fig. S3.**
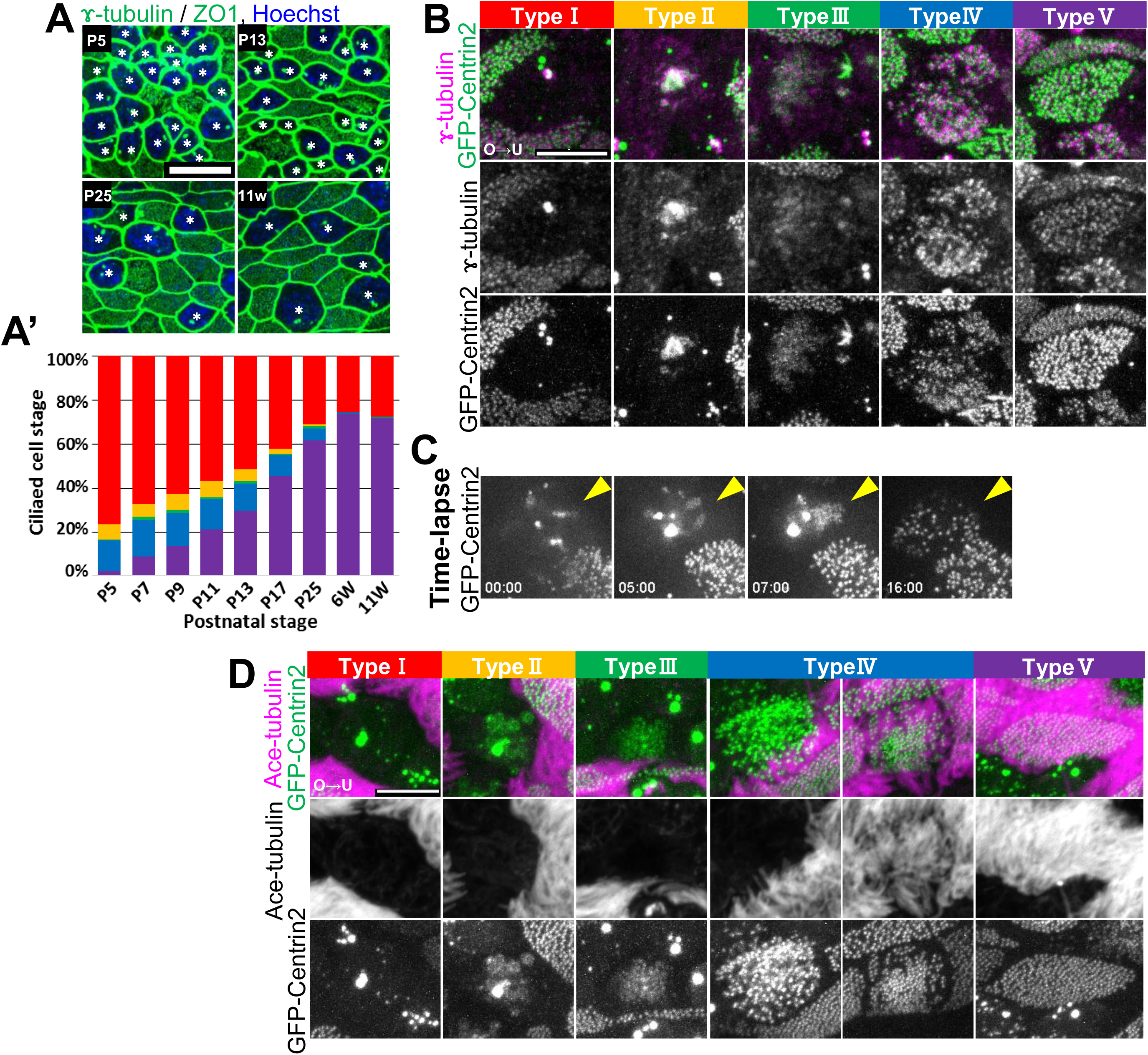
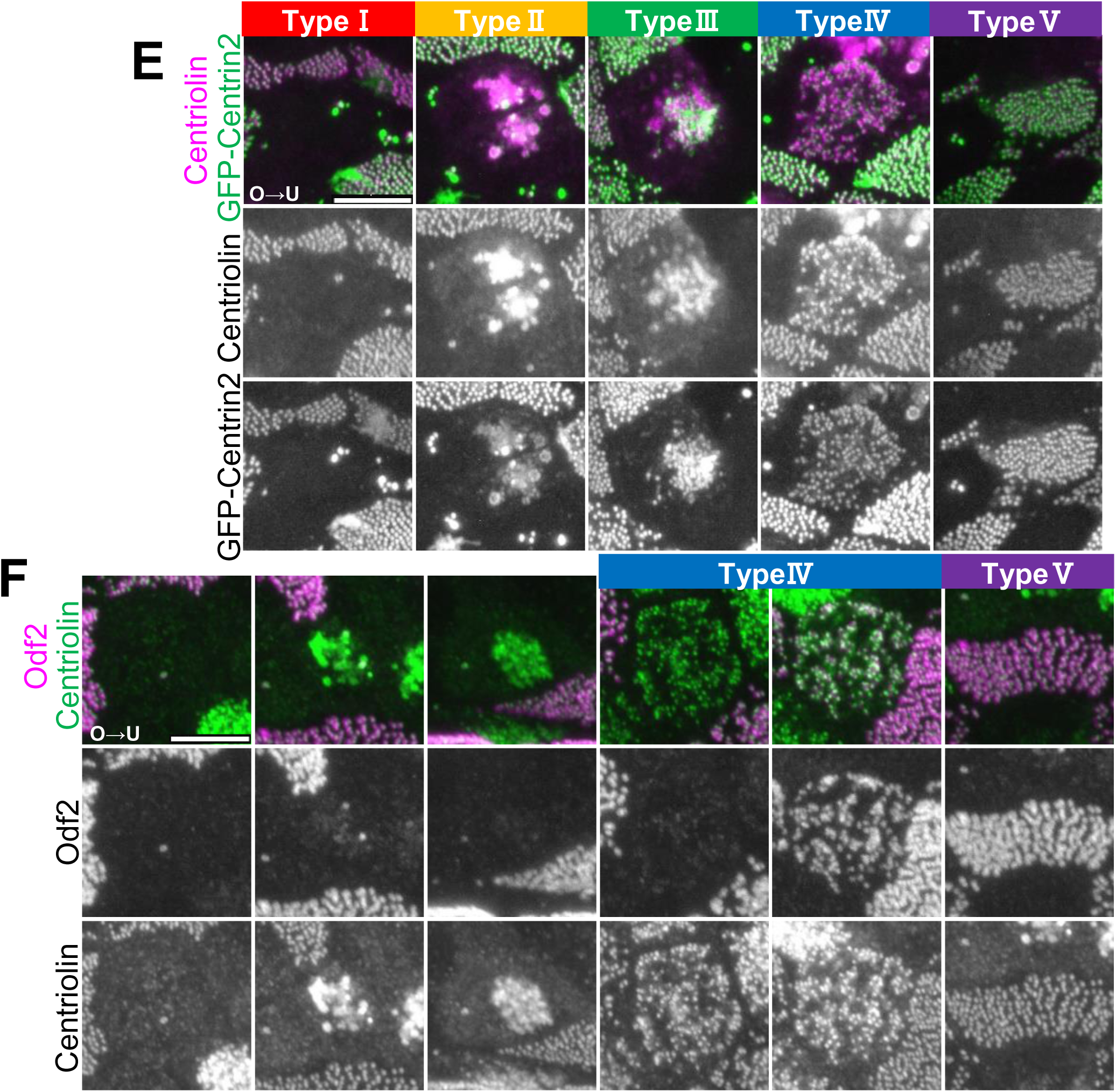
Cell type transition of MCCs and localization of BB proteins during the differentiation process of MCCs. A, Representative images of oviducts on postnatal day 5 (P5) to 11 weeks (11w) stained with anti-ZO1 (green), anti–γ-tubulin (green), and Hoechst (blue). Secreting cells (which are categorized as type I) are shown with asterisks. (Bar = 10 μm) A’, Changes in cell type composition at each developmental stage. Oviducts from three animals were used for the analysis. Analyzed cell numbers; 1469 (P5), 2341 (P7), 1999 (P9), 1609 (P11), 1791 (P13), 2077 (P17), 2807 (P25), 2493 (6w), and 2870 (11w). B, Confocal images of *GFP-CETN2* (green) oviduct stained with anti–γ-tubulin (magenta). MIP images were rendered from 30 images acquired at a step size of 0.2 μm. C, Snapshot images after time-lapse recording (same as Sup. Mov. 2) of *GFP-CETN2*. Arrowheads indicate the cell of interest undergoing cell type transition. Timepoints are shown on the bottom. D, Confocal images of the oviduct of P13 *GFP-CETN2* (green) stained with anti-acetylated tubulin antibody (magenta). MIP images rendered from 50 slices acquired at a step size of 0.2 μm. (Bar = 5 μm) E, Confocal images of the oviduct of P13 *GFP-CETN2* (green) stained with anti-Centriolin antibody (magenta). MIP images rendered from 30 slices acquired at a step size of 0.2 μm. To compare signal intensities between cell types, all the images were cropped after simultaneously acquired images. F, Confocal images of P13 WT oviduct stained with anti-Centriolin (green) and anti-Odf2 (magenta) antibodies. MIP images rendered from 30 slices acquired at a step size of 0.2 μm. All the images were cropped after from simultaneous acquisition. Two representative type IV cells with different Odf2 staining patterns are shown. Odf2 localized to BBs in 83% (183/221) of type IV cells and 100% of type V cells.

**Fig. S4.**
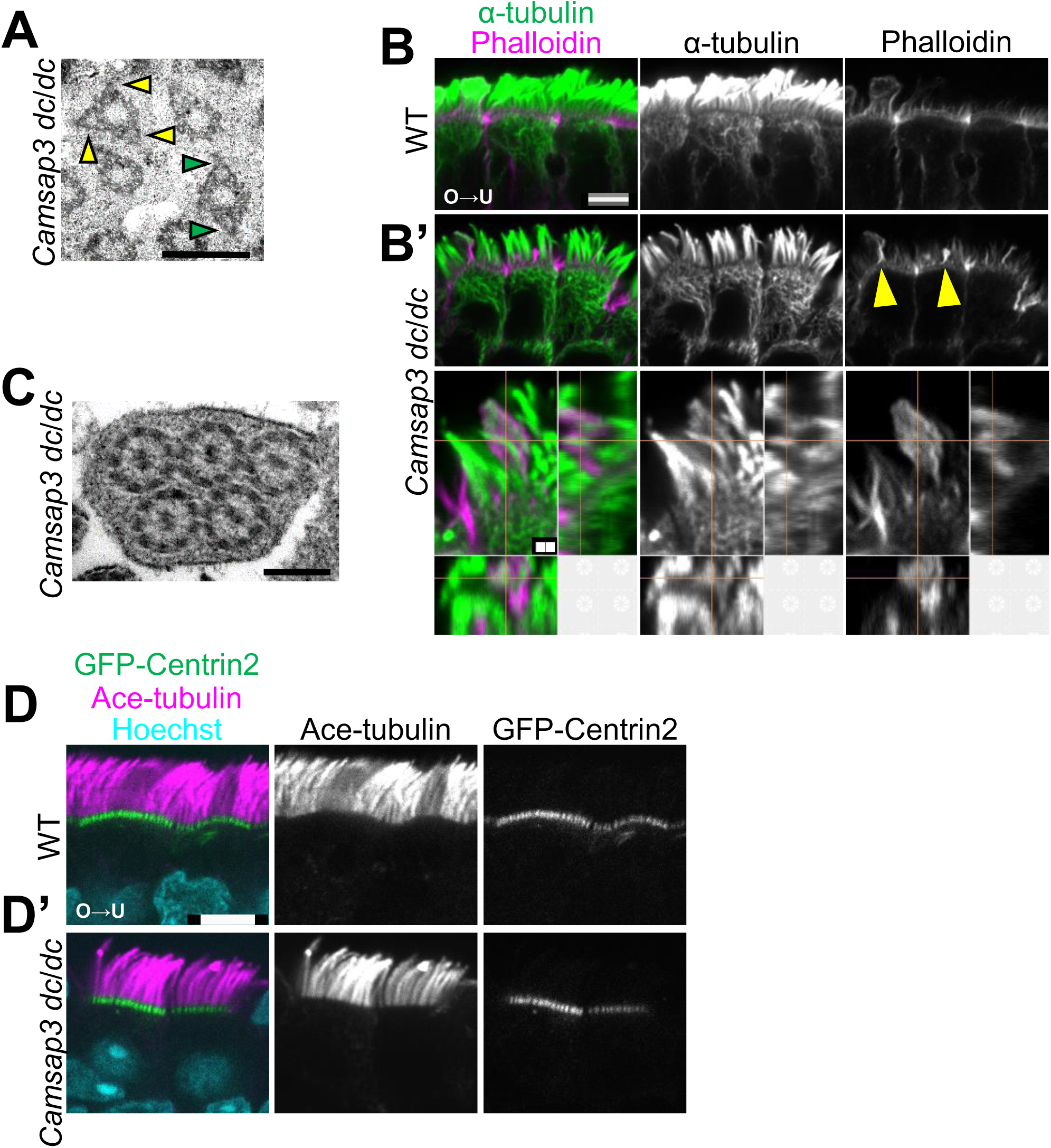
Distribution of microtubules in *Camsap3 dc/dc* mutant MCCs. A, Images of BBs observed with a transmission electron microscopy showing examples of abnormal BBs with multiple protrusions resembling BF (shown with arrow heads). (Bar = 0.5 μm) B and B’, Lateral view of oviducts of WT (B) and *Camsap3 dc/dc* (B’) stained with anti–α-tubulin antibody (green) and phalloidin (magenta). Bottom rows of (B’) show 3D orthogonal views of abnormal protrusions with phalloidin signals (indicated with arrowheads). (Bar = 5 μm (top) and 1 μm (bottom)) C, Electron micrograph of abnormal cilia. Multiple axonemes are covered within the cell membrane. (Bar = 0.2 μm) D and D’, Lateral view of MCCs of WT and *Camsap3 dc/dc* animals with *GFP-CETN2* (green) background stained with anti-acetylated tubulin antibody (magenta) and Hoechst (cyan). (Bar = 5 μm)

**Fig. S5.**
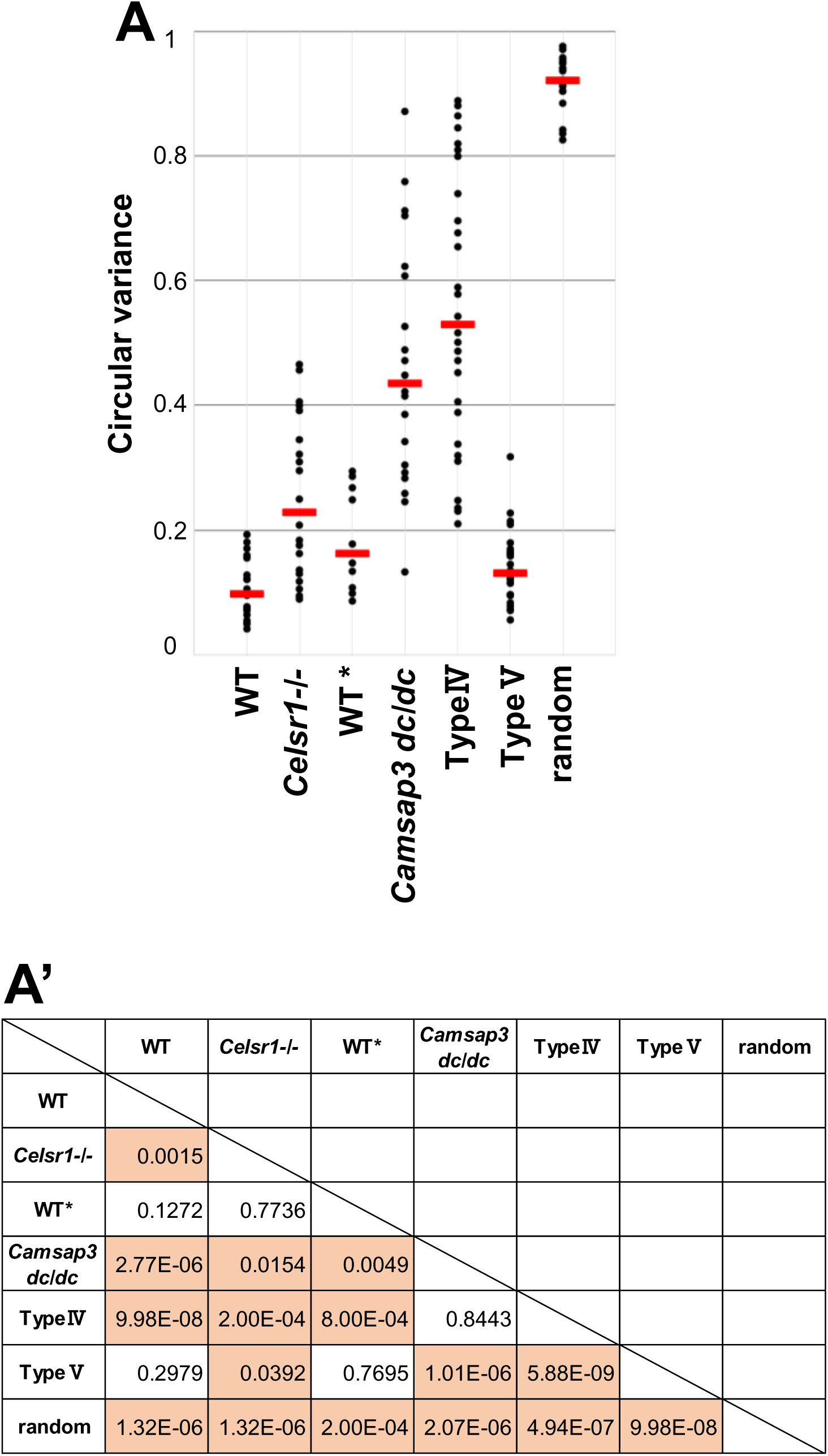
Statistical comparison of CVs between genotypes and cell types. A, Data sets from Fig. 1D, 4E, and 5C were summarized for comparison. The distribution of CVs is shown. Red lines represent the median CV. “WT” and “*Celsr1-/-*” show results of the 12 w old (same data as shown in Fig. 1D, n = 20 cells). “WT*” and “*Camsap3 dc/dc,*” results of adult littermates (shown in Fig. 5C, n = 10 and 20 cells in WT and *Camsap3 dc/dc,* respectively). “Type IV” and “Type V,” results of WT P13 (shown in Fig. 4E, n= 28 cells in each cell type). “random” is the result of a calculation using random numbers corresponding to 150 BBs, n = 20 cells. B, A table showing P values after the comparison between two groups. Non-parametric multiple comparisons were performed using the Steel-Dwass method. *p* values smaller than 0.05 are shown in orange.

**Fig. S6.**
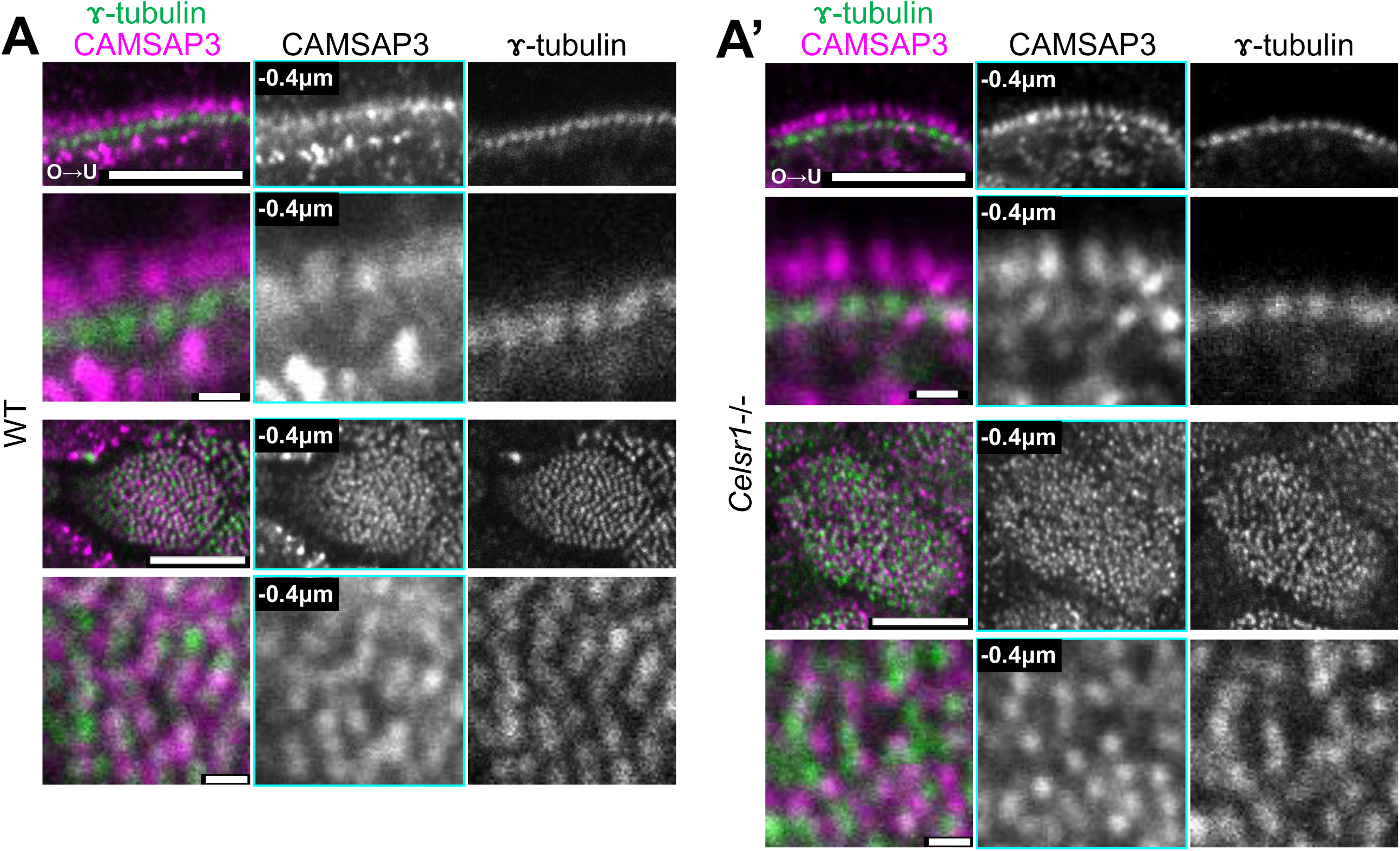
CAMSAP3 localization to the base of cilia is maintained in *Celsr1*-/- mutant. A and A’, WT and *Celsr1*-/- mutant (adult) MCCs stained with anti-CAMSAP3 (magenta) and anti–γ-tubulin (green) antibodies; lateral view (top rows) and apical view (bottom rows). All images were single plane. The focal plane of CAMSAP3 was 0.4 μm apical to the focal plane of γ-tubulin, and is indicated as “-0.4 μm” (cyan frame). Bottom panels show the magnified images. (Bar = 5 μm (top) and 0.5 μm (bottom))

Key resource table, Antibodies used in this study with fixation conditions

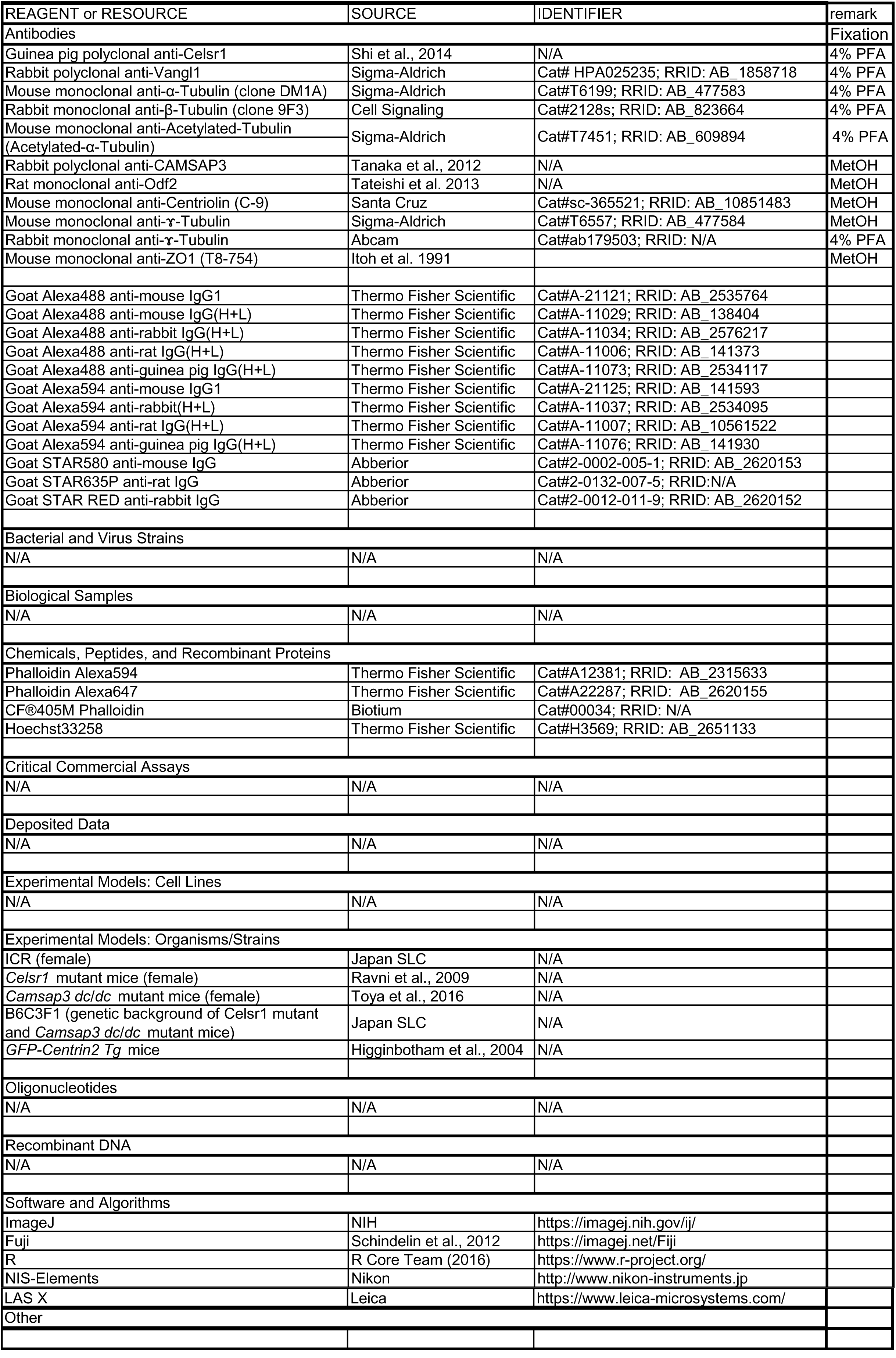

**Supplementary movie 1, Confocal microscopy images of tubulin in a MCC along the apical-basal axis**

GFP-Centrin2 (green) and β-tubulin (magenta) in an oviduct MCC (the same cell shown in Fig. 2D). From apical to basal, the *z*-step size (0.2 μm) is shown on the bottom.

**Supplementary movie 2, Time-lapse observation of MCC maturation**

Time-lapse recording of a cell expressing GFP-Centrin2. Snapshots are shown in Fig. S3C. The timepoint is indicated at the bottom (in hours). MIP images were rendered from micrographs acquired at a step size of 0.5 μm.

